# Direct detection of endogenous Gαi activity in cells with a sensitive conformational biosensor

**DOI:** 10.1101/2024.08.21.609006

**Authors:** Alex Luebbers, Remi Janicot, Jingyi Zhao, Clementine E. Philibert, Mikel Garcia-Marcos

**Affiliations:** Department of Biochemistry & Cell Biology, Chobanian & Avedisian School of Medicine, Boston University, Boston, MA 02118, USA; Department of Biology, College of Arts & Sciences, Boston University, Boston, MA 02115, USA

## Abstract

Activation of heterotrimeric G-proteins (Gαβγ) by G-protein-coupled receptors (GPCRs) is not only a mechanism broadly used by eukaryotes to transduce signals across the plasma membrane, but also the target for a large fraction of clinical drugs. However, approaches typically used to assess this signaling mechanism by directly measuring G-protein activity, like optical biosensors, suffer from limitations. On one hand, many of these biosensors require expression of exogenous GPCRs and/or G-proteins, compromising readout fidelity. On the other hand, biosensors that measure endogenous signaling may still interfere with the signaling process under investigation or suffer from having a small dynamic range of detection, hindering broad applicability. Here, we developed an optical biosensor that detects the endogenous G-protein active species Gαi-GTP upon stimulation of endogenous GPCRs more robustly than current state-of-the-art sensors for the same purpose. Its design is based on the principle of bystander Bioluminescence Resonance Energy Transfer (BRET) and leverages the Gαi-binding protein named GINIP as a high affinity and specific detector module of the GTP-bound conformation of Gαi. We optimized this design to prevent interference with G_i_-dependent signaling (cAMP inhibition) and to enable implementation in different experimental systems with endogenous GPCRs, including neurotransmitter receptors in primary astroglial cells or opioid receptors in cell lines, which revealed opioid neuropeptide-mediated activation profiles different from those observed with other biosensors involving exogenous GPCRs and G-proteins. Overall, we introduce a biosensor that directly and sensitively detects endogenous activation of G-proteins by GPCRs across different experimental settings without interfering with the subsequent propagation of signaling.

## INTRODUCTION

Heterotrimeric G proteins (Gαβγ) are quintessential mediators of intercellular communication (*1–4*). Defining the molecular mechanisms by which they are regulated is of paramount importance because they impact a vast range of physiological processes and diseases. This is well exemplified by the ongoing interest in G protein-coupled receptors (GPCRs), which are the canonical activators of G proteins. GPCRs are receptors displayed at the cell surface that, upon stimulation, activate an intracellular G protein, a transducer that leads to a cellular response (*1, 3, 5, 6*). This evolutionarily conserved mechanism of signal transduction is very versatile, as it instructs intracellular responses to numerous extracellular stimuli of diverse nature, including neurotransmitters, hormones, light, odorants, or mechanical cues, among others (*3, 5, 7–9*). The medical relevance of GPCRs is evident not only because they serve as pharmacological targets for >30% of clinically approved drugs, but also because they remain actively pursued for the development of new and improved therapeutic approaches (*10–12*). For example, opioid drugs exert their potent analgesic effects by targeting the same GPCRs that are activated by endogenous neuropeptides like endorphins, enkephalins, or dynorphins (*13*). These GPCRs, including the μ-opioid receptor (MOR) and the δ-opioid receptor (DOR) among others, have been the subject of intense pharmacological research to develop safer analgesic drug alternatives with reduced deleterious side-effects like dependency or respiratory depression, although some other GPCRs have also started to emerge as potential targets for this purpose (*14–19*).

Mechanistically, heterotrimeric G protein signaling starts with GPCRs acting as Guanine nucleotide exchange factors (GEFs) — i.e., promoting the exchange of GDP for GTP in Gα subunits, leading to formation of Gα-GTP and free Gβγ species that modulate downstream effectors (e.g., adenylyl cyclases) to propagate signaling. Based on the structural and functional similarities of Gα subunits, G proteins are classified into four families: G_i/o_, G_s_, G_q/11_, or G_12/13_ (*1*). The specificity of GPCRs for coupling to different G proteins displays varying degrees of selectivity; some GPCRs recognize a particular family of G proteins with high specificity, whereas other GPCRs couple promiscuously to G proteins across different families (*20*). The identity of the G protein dictates the nature of the cellular response elicited by acting on specific downstream effectors. For example, Gαs-GTP formed upon activation of the β2 adrenergic receptor (β2AR) stimulates the effector adenylyl cyclase, whereas Gαi-GTP formed upon activation of the GABA_B_ receptor (*21*) or opioid receptors (*22–24*) inhibits it. These opposing actions translate into the corresponding effects on the cellular levels of the second messenger cAMP synthesized by adenylyl cyclases, which dictates various cell responses.

A general strategy to measure G protein signaling responses is to use indirect approaches, including the measurement of downstream second messengers like cAMP. Another general strategy is to directly measure the formation of active G protein species, which is frequently done using optical biosensors based on resonance energy transfer (RET) methods with fluorescent or bioluminescent donors (FRET or BRET, respectively) (*25–29*). Indirect approaches are subject to crosstalk or signal amplification events that compromise the fidelity of the readout as a representation of the GPCR-G protein signal transduction event. While biosensors that directly measure G protein activation in real time greatly alleviate these issues, they are not devoid of limitations. For example, a broad class of biosensor designs that monitor the dissociation of Gα and Gβγ subunits (*26–28, 30*) requires the expression of multiple genetic components including exogenous, tagged G proteins. This has two potentially undesired consequences. One is that overexpression of exogenous G proteins might distort the readout and interfere with endogenous GPCR signaling (*31*). The other consequence is that the need for simultaneous expression of multiple genetic components restricts their implementation to easily transfectable cell lines. The latter scenario in cell lines also tends to be accompanied by the need to express exogenous GPCRs to detect responses, which skews the system further away from a native cellular condition. Thus, these widely adopted biosensors are not well suited to investigate endogenous GPCR activity, especially in physiologically-relevant systems like primary cells.

More recently, another broad class of biosensors has been developed to detect Gα-GTP instead of Gα/Gβγ dissociation, which have overcome some of the limitations in terms of preservation of signaling fidelity and of applicability across physiologically relevant systems. The first example of this class of biosensors was a platform based on the <u>B</u>RET biosensor with <u>ER/K</u> linker and <u>Y</u>FP (BERKY) design (*32*). These biosensors consist of a single polypeptide chain that permits the detection of endogenous Gα-GTP generated by endogenous GPCRs in different experimental settings, including primary cells like neurons, and without interfering with GPCR-G protein signaling to downstream signaling targets (*32*). While BERKY biosensors overcome many of the limitations of preceding biosensor designs, the modest dynamic range of detection for endogenous responses has probably hindered their wider applicability. Other biosensor platforms to detect Gα-GTP developed subsequently, like <u>ONE</u> vector <u>G</u> protein <u>O</u>ptical (ONE-GO, (*33*)) biosensors, or the <u>E</u>ffector <u>M</u>embrane <u>T</u>ranslocation <u>A</u>ssay (EMTA, (*34*)), have improved the dynamic range of detection of G protein activation by endogenous GPCRs, albeit at the expense of other limitations. For example, the ONE-GO sensors design is based on assembling and delivering a multicomponent biosensor system with a single vector, allowing for the measurement of responses triggered by endogenous GPCRs in a remarkably wide range of primary cell types and without interfering with downstream signaling (*33*), yet it requires expression of trace amounts of exogenous, tagged Gα subunits. As for the EMTA system, even though it was shown to work with endogenous, untagged Gα subunits for some types of G proteins (*34*), its applicability for endogenous GPCRs across physiologically-relevant systems like primary cells has not been established yet. The latter may be related to the difficulty of delivering the multiple genetic components composing this type of sensor to cells. Furthermore, it is likely that EMTA components interfere with GPCR signaling, a potential limitation that has not been assessed yet (*34*). For example, EMTA biosensors for G proteins of the G_i/o_ family are based on using Rap1GAP as a detector module, for which it is unclear whether it affects nucleotide exchange on different Gα subunits of this family or its preference for binding to Gα-GTP or Gα-GDP dissociated from Gβγ (*35, 36*), thereby raising questions about what is exactly represented by the BRET changes detected by this sensor. Thus, there is still a critical unmet need to develop biosensors for the detection of endogenous G protein activity that combine a large dynamic range of detection with the lack of interference with GPCR signaling and potential for broad applicability across experimental settings.

Here, we introduce a BRET biosensor design that detects endogenous Gαi-GTP, even when produced upon stimulation of endogenous GPCRs in cell lines or primary cells, without interfering with signaling to downstream effectors. We focused on Gαi to develop the new design based on the availability of a recently characterized Gαi-binding protein, GINIP, which was leveraged as a critical component of the biosensor to sensitively and specifically detect the active conformation of the G protein. We also optimized an initial prototype to abolish interference with signaling and to facilitate implementation in different experimental systems by assembling all sensor components in a single vector. We showcase the versatility of this biosensor design by implementing it in a broad range of formats, from transient transfection to generation of stable cell lines to short-term lentiviral transduction of primary cells, and by demonstrating its utility in characterizing responses from many different GPCRs and many different ligands, including the profiling of the activity of opioid neuropeptides on endogenously expressed opioid receptors.

## RESULTS

### Detection of endogenous Gαi-GTP in cells via bystander BRET

We envisioned a bioluminescence resonance energy transfer (BRET)-based biosensor design for the detection of endogenous Gαi activation based on two components: a detector module for Gαi-GTP fused to the BRET donor nanoluciferase (Nluc), and a membrane-anchored BRET acceptor fluorescent protein (YFP) (**Fig. 1A**). The principle of this design is that the BRET donor would be recruited from the cytosol to the plasma membrane upon activation of membrane-resident Gαi subunits, which would in turn lead to BRET with the acceptor due to the increased proximity and crowding effects on the two-dimensional plane of the membrane— i.e., a phenomenon known as bystander BRET (*37, 38*). We reasoned that the recently characterized GPCR signaling modulator GINIP would serve as a module to detect active Gαi with high sensitivity and specificity based on its high affinity (K_D_∼65 nM) for the G protein in its GTP-bound conformation but not is its GDP-bound one (*39, 40*). GINIP binds similarly to the three Gαi isoforms, Gαi1, Gαi2 and Gαi3, but not to other G proteins of the G_i/o_ family like Gαo and Gαz, or to members of other G protein families (*39*). GINIP does not affect directly nucleotide binding or hydrolysis by Gα (*39*), which we reasoned would minimize the potential interference of our biosensor design with the signaling process to be measured. To test this design, we co-expressed GINIP-Nluc and YFP-CAAX (*38*) (a fusion of YFP and the C-terminal sequence of KRas containing a polybasic sequence and prenylation box for plasma membrane targeting) with the GABA_B_ receptor (GABA_B_R) in HEK293T cells (**Fig. 1B, C**). No exogenous G protein was expressed. Stimulation of the GABA_B_R led to a marked increase in BRET that was rapidly reverted upon addition of an antagonist. This response was efficiently suppressed by pertussis toxin (PTX) (**Fig. 1B**) or by a mutation in GINIP (W139A) that disrupts its binding to G proteins (*40*) (**Fig. 1C**), indicating that the BRET change represents Gαi activity. Using this biosensor, we obtained concentration-response curves not only for GABA_B_R, but also for three other G_i_-coupled GPCRs: α2_A_-adrenergic receptor (α2_A_-AR), dopamine 2 receptor (D2R), and μ-opioid receptor (MOR). These results demonstrate that, when co-expressed in cells, GINIP-Nluc and YFP-CAAX constitute a bystander BRET sensor for endogenous Gαi activation downstream of multiple GPCRs.

**Figure 1.**
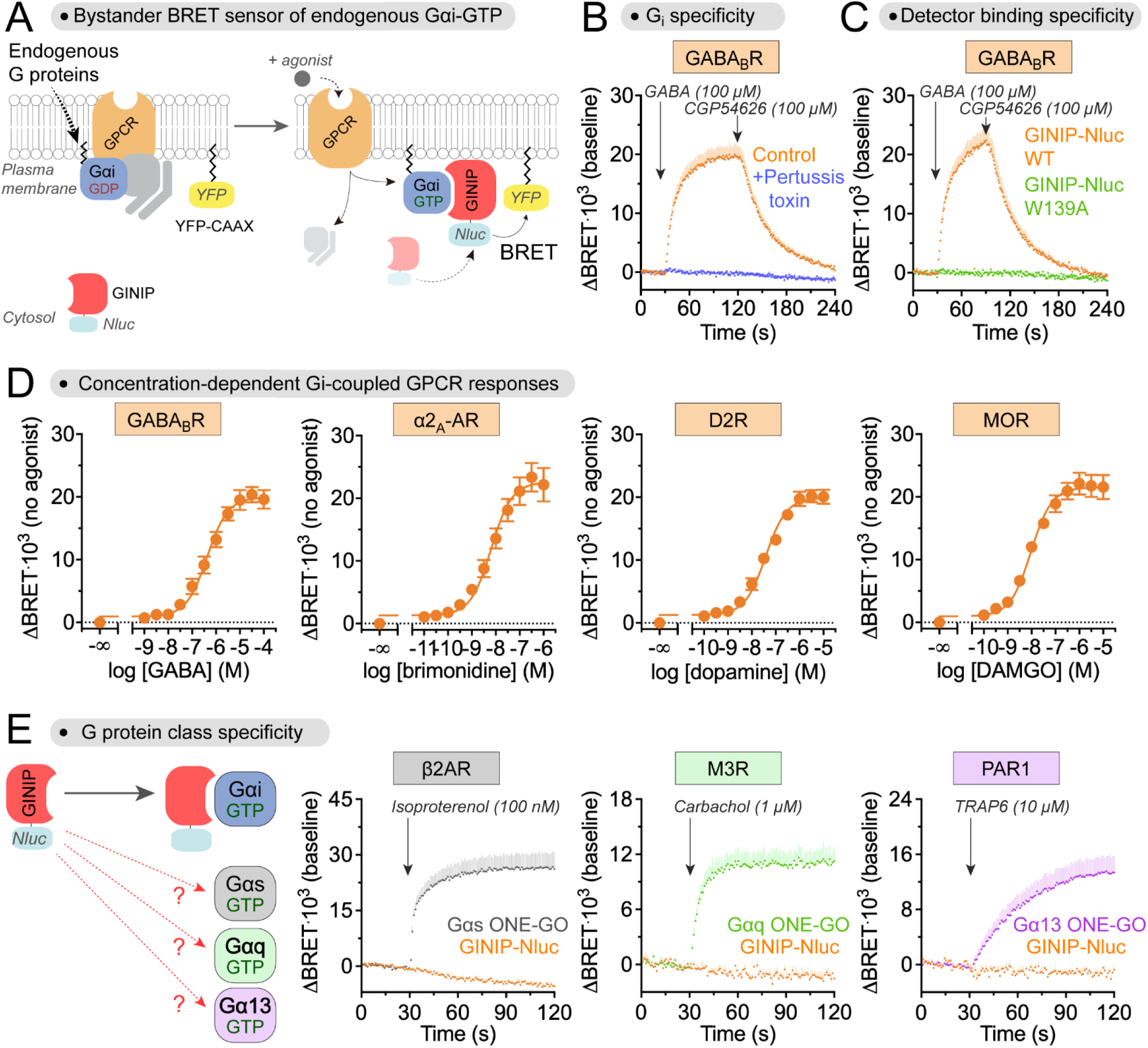
Detection of endogenous Gαi-GTP with a bystander BRET biosensor based on GINIP. **(A)** Diagram showing the detection of endogenous Gαi-GTP via bystander BRET when nanoluciferase (Nluc)-fused GINIP (BRET donor) in the cytosol is recruited to the proximity of membrane anchored YFP (YFP-CAAX, BRET acceptor) due to binding to membrane-bound Gαi-GTP. **(B)** Responses detected by GINIP-based bystander BRET sensor depend on GPCR-mediated activation of G_i_. Kinetic BRET measurements were carried out in HEK293T cells expressing GABA_B_R, GINIP-Nluc, and YFP-CAAX (but no exogenous G protein) in the absence (orange) or presence (blue) of Pertussis toxin via PTX-S1 expression. Cells were treated with GABA and CGP54626 as indicated. Mean ± S.E.M., n=4. **(C)** Gαi-GTP bystander BRET sensor relies on the interaction between GINIP and Gαi. Kinetic BRET measurements were carried out as in (B), except that GINIP-Nluc WT (orange) was compared to cells expressing a GINIP-Nluc construct bearing the G protein binding-deficient mutant W139A (green). Mean ± S.E.M., n=3 for GINIP-Nluc WT, and n=2 for GINIP-Nluc W139A. **(D)** Gαi-GTP bystander BRET sensor detects responses to multiple G_i_-coupled GPCRs. BRET was measured in HEK293T cells expressing GINIP-Nluc WT and YFP-CAAX along with the indicated GPCRs upon stimulation with their cognate agonists. Mean ± S.E.M., n=3. **(E)** Gαi-GTP bystander BRET sensor does not detect activation of G_s_, G_q_, or G_13_. BRET was measured in HEK293T cells expressing GINIP-Nluc WT and YFP-CAAX (orange) along with the indicated GPCRs upon stimulation with their cognate agonists. In parallel experiments, BRET was measured in HEK293T cells expressing Gαs ONE-GO (grey), Gαq ONE-GO (green), and Gα13 ONE-GO (magenta) along with the indicated GPCRs upon stimulation with their cognate agonists. Mean ± S.E.M., n=3.

### Detection of Gαi-GTP with the GINIP-based bystander sensor displays a large dynamic range

To benchmark the performance of the newly developed bystander BRET sensor, we compared it to a current “gold standard” for the direct detection of endogenous G protein activity— i.e., BERKY biosensors (*32*). When compared side by side with the BERKY biosensor for Gαi-GTP (i.e., Gαi*-BERKY3), the newly developed GINIP-based bystander BRET sensor led to much larger responses (∼10-fold) upon stimulation of GABA_B_R in HEK293T cells expressing exclusively endogenous G proteins (**Fig. S1**). This indicates that the bystander BRET sensor outperforms previously described BERKY biosensors for the detection of endogenous Gαi-GTP, leading to an improvement in the dynamic range of the responses detected.

### GINIP-based bystander Gαi sensor does not detect the activation of G proteins of other families

Next, we assessed the selectivity of the bystander BRET sensor for detecting Gαi over other types of G proteins. For this, we tested whether the sensor would detect responses upon stimulation of GPCRs that activate representative members of the other families of G proteins (G_s_, G_q/11_, and G_12/13_, instead of G_i/o_) (**Fig. 1E**), with the expectation that they would not because GINIP only binds to Gαi1, Gαi2 and Gαi3 (*39, 41*). Stimulation of the β2 adrenergic receptor (β2AR), the M3 muscarinic acetylcholine receptor (M3R), or the protease-activated receptor 1 (PAR1), which are known to activate G_s_, G_q/11_, or G_12/13_ (*20, 34*), respectively, did not lead to a BRET response in HEK293T cells expressing GINIP-Nluc and YFP-CAAX (**Fig. 1E**). This was not because of lack of activation of the cognate G proteins, as we detected their activation using another type of biosensor (i.e., ONE-GO, (*33*)) in parallel experiments with the same GPCRs (**Fig. 1E**). These observations validate that the bystander BRET sensor specifically detects Gαi activity without contribution of G proteins of other families to the observed responses.

### Gαi bystander BRET sensor moderately affects cAMP regulation in cells

After establishing the specificity of the Gαi bystander BRET sensor, we set out to determine if its expression interfered with G protein signaling to downstream targets in cells, such as inhibition of adenylyl cyclase. While GINIP does not affect the ability of Gαi to bind or hydrolyze nucleotides, it can block Gαi binding to adenylyl cyclase when expressed at sufficiently high levels (*39*). To test the potential effect of GINIP-Nluc expression on adenylyl cyclase regulation, we measured cAMP levels in cells upon GPCR stimulation using Glo-Sensor, a luminescence-based probe (*42*). More specifically, we measured the inhibition of isoproterenol-elicited cAMP by GABA_B_R-activated G_i_ in the presence or absence of Gαi bystander BRET sensor (**Fig. 2A**). While expression of the sensor under the same conditions as in experiments to detect endogenous G_i_ responses did not affect the maximal inhibition achieved upon GABA_B_R stimulation (**Fig. 2A**, *right*) or the expression of G proteins (**Fig. 2B**, *right*), it modestly decreased the potency of the inhibition by GABA (∼4-fold increase in the IC_50_) (**Fig. 2B**, *left*). These results suggest that the Gαi bystander BRET sensor has modest, yet detectable, effects on cellular responses mediated by G_i_ proteins upon GPCR stimulation.

**Figure 2.**
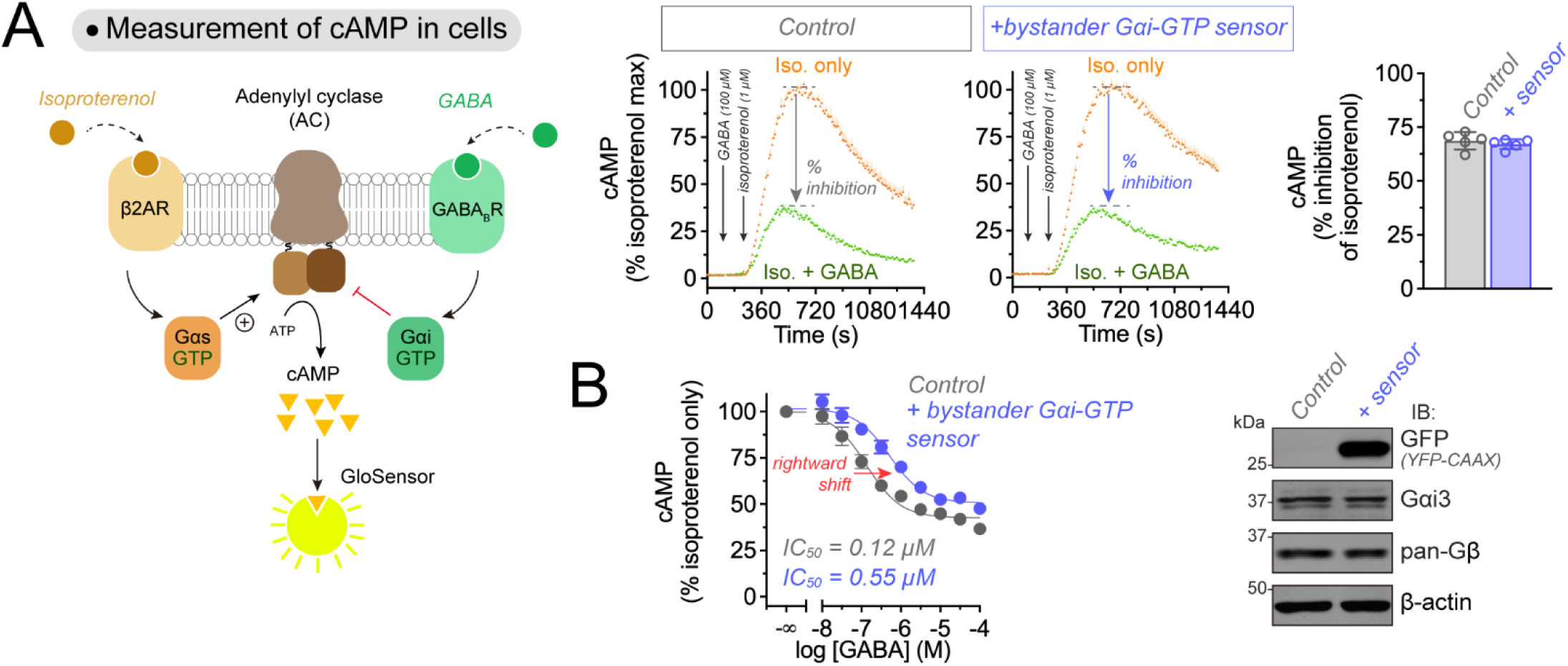
Effect of Gαi-GTP bystander BRET sensor on G_i_-mediated inhibition of G_s_-stimulated adenylyl cyclase activity. **(A)** *Left,* Diagram showing GPCR-G protein mediated regulation of adenylyl cyclase (AC) activity and subsequent detection of cAMP levels in cells via a luminescence-based biosensor (GloSensor). *Right,* Gαi-GTP bystander BRET sensor does not affect the efficacy of G_i_-mediated inhibition of AC activity. Kinetic luminescence measurements of cAMP levels in HEK293T cells were carried out in the absence (grey) or presence (blue) of Gαi-GTP bystander BRET sensor expression. Cells were treated with isoproterenol with (green) or without (orange) pretreatment with GABA. The percentage of GABA-mediated inhibition of the isoproterenol response is quantified on the graph on the right. Mean ± S.E.M., n=5. **(B)** *Left,* Gαi-GTP bystander BRET sensor modestly reduces the potency of G_i_-mediated inhibition of AC. Concentration-dependent measurements of cAMP inhibition by GABA were carried out in the absence (grey) or presence (blue) of Gαi-GTP bystander BRET sensor expressions. Cells were stimulated with isoproterenol in the presence of the indicated concentrations of GABA. Mean ± S.E.M., n=3. *Right,* A representative immunoblotting result confirms the expression of the bystander sensor and that it does not affect expression of endogenous G proteins.

### Detection of endogenous Gαi-GTP with a single-vector system for biosensor expression

We set out to minimize or completely eliminate the interference of the bystander BRET sensor with G_i_ signaling. For this, we took inspiration from the recently described ONE-GO biosensor design (*33*). This design allows for sensitive detection of G protein activity without interfering with it by virtue of expression of the sensor components at reduced levels, yet at relative ratios adequate for the detection of large BRET differences (*33*). We mimicked the ONE-GO design by expressing the GINIP-Nluc cassette after a low efficiency IRES (IRES*) downstream of the YFP-CAAX component, which was placed right after the promoter, with the overall intent of favoring higher acceptor-to-donor expression ratios to maximize the magnitude of BRET differences (**Fig. 3A**). The construct was assembled in a plasmid backbone suitable for lentiviral packaging to facilitate its potential application in cell types not easily transfected. This design was named “bONE-GO biosensor”, for <u>b</u>ystander <u>ONE</u> vector <u>G</u> protein <u>O</u>ptical biosensor (**Fig. 3A**). We reasoned that reduced expression of GINIP-Nluc would (1) reduce the potential interference with G_i_ signaling, and (2) help achieving a high acceptor-to-donor ratio conducive to adequate detection of BRET differences. HEK293T cells expressing the bONE-GO sensor and GABA_B_R, but no exogenous G protein, elicited a robust BRET response upon GABA stimulation, which was rapidly reverted upon application of a GABA_B_R antagonist (**Fig. 3B**). This BRET response was suppressed by pertussis toxin, indicating that is was dependent on activation of G_i_ (**Fig. 3B**). We obtained concentration-response curves for GABA_B_R and three other G_i_-coupled GPCRs, α2_A_-AR, D2R, and MOR (**Fig. 3C**). These results indicate that, much like its multi-plasmid predecessor, the bONE-GO design detects endogenous Gαi-GTP levels and is broadly applicable across receptors that activate G_i_.

**Figure 3.**
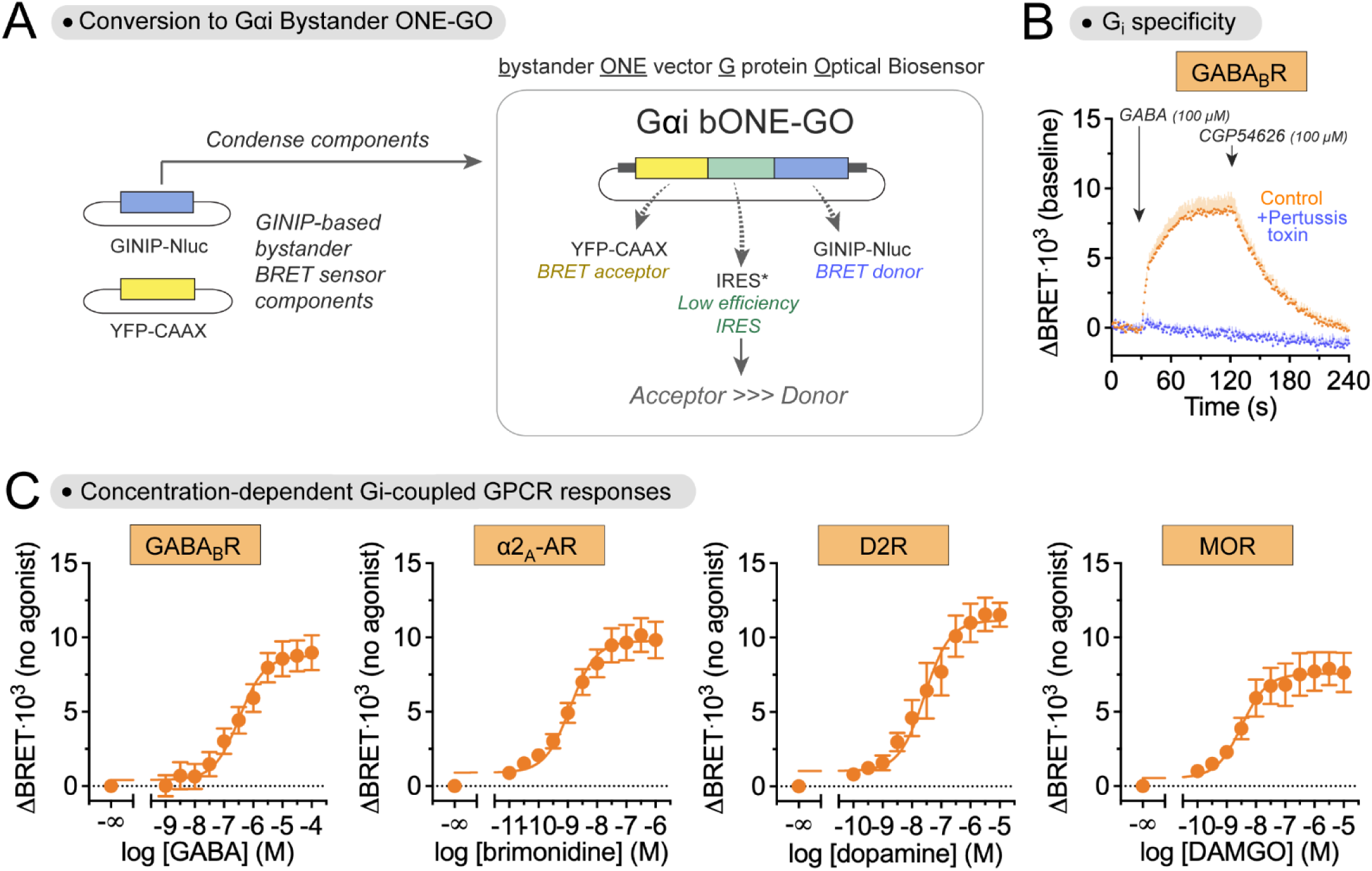
Generation of a Gαi bystander ONE vector G protein Optical (Gαi bONE-GO) for detecting endogenous Gαi-GTP. **(A)** Diagram showing the conversion of the multi-plasmid Gαi-GTP bystander BRET sensor to the single-plasmid Gαi bystander ONE vector G protein Optical Biosensor (bONE-GO). **(B)** Responses detected by Gαi bONE-GO sensor depend on GPCR-mediated activation of G_i_. Kinetic BRET measurements were carried out in HEK293T cells expressing GABA_B_R and Gαi bONE-GO (but no exogenous G protein) in the absence (orange) or presence (blue) of Pertussis toxin via PTX-S1 expression. Cells were treated with GABA and CGP54626 as indicated. Mean ± S.E.M., n=4. **(C)** Gαi bONE-GO sensor detects responses to multiple G_i_-coupled GPCRs. BRET was measured in HEK293T cells expressing Gαi bONE-GO along with the indicated GPCRs upon stimulation with their cognate agonists. Mean ± S.E.M., n=4 (for GABA_B_R, α2_A_-AR, MOR), n=3 (for D2R).

### Gαi bONE-GO sensor does not affect cAMP regulation

Having established that Gαi bONE-GO detects endogenous responses, we set out to test whether it interfered with G_i_-mediated signaling. Mirroring the experiments performed in **Fig. 2** with its multi-plasmid predecessor, we assessed the changes in GPCR-modulated cAMP levels in cells expressing Gαi bONE-GO compared to controls (**Fig. 4A**). We found that expression of Gαi bONE-GO under the same conditions as in experiments detecting endogenous Gαi-GTP (e.g., **Fig. 3**), did not affect either the efficacy (i.e., maximal effect) or potency (i.e., IC_50_) of GABA_B_R-mediated inhibition of isoproterenol-elicited cAMP responses (**Fig. 4A, B**). Protein levels of Gαi3 or Gβ were also not affected by Gαi bONE-GO expression (**Fig. 4B**). The GINIP-Nluc module of the biosensor was undetectable by immunoblotting (*not shown*), and the YFP-CAAX module was barely detectable (**Fig. 4B**). Since GINIP-Nluc is expressed after a low efficacy IRES in Gαi bONE-GO, its expression must be exceedingly low, thereby explaining why it does not affect G_i_ signaling in cells. In summary, the bONE-GO design allows for sensitive detection of endogenous Gαi activation without interfering with the propagation of signaling from the G protein to downstream effectors.

**Figure 4.**
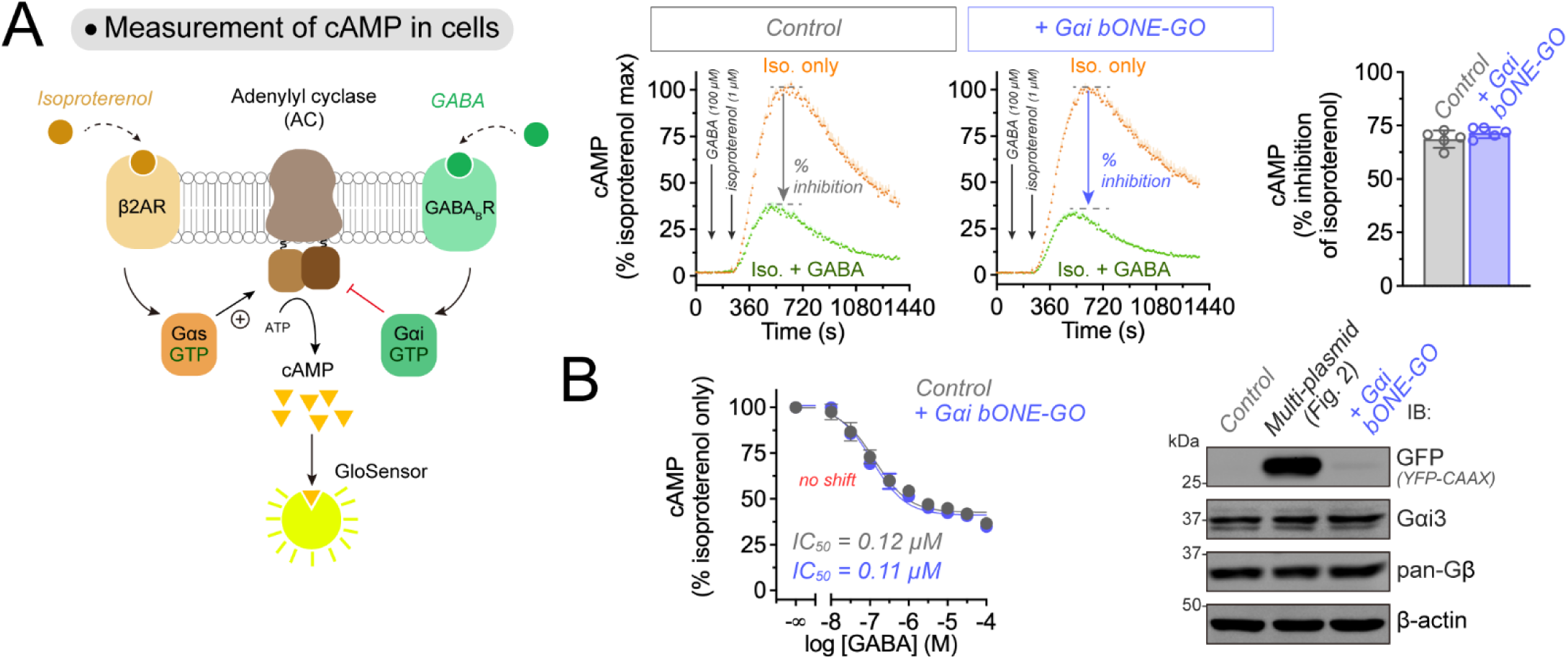
Effect of Gαi bONE-GO on G_i_-mediated inhibition of G_s_-stimulated adenylyl cyclase activity. **(A)** *Left,* Diagram showing GPCR-G protein mediated regulation of adenylyl cyclase (AC) activity and subsequent detection of cAMP levels in cells via a luminescence-based biosensor (GloSensor). *Right,* Gαi-GTP bONE-GO sensor does not affect the efficacy of G_i_-mediated inhibition of AC activity. Kinetic luminescence measurements of cAMP levels in HEK293T cells were carried out in the absence (grey) or presence (blue) of Gαi bONE-GO sensor expression. Cells were treated with isoproterenol with (green) or without (orange) pretreatment with GABA. The percentage of GABA-mediated inhibition of the isoproterenol response is quantified on the graph on the right. Mean ± S.E.M., n=5. Data for the “Control” condition are the same as for the “Control” presented in Figure 2. **(B)** *Left,* Gαi-GTP bONE-GO sensor does not affect the potency of G_i_-mediated inhibition of AC. Concentration-dependent measurements of cAMP inhibition by GABA were carried out in the absence (grey) or presence (blue) of Gαi-GTP bONE-GO sensor expressions. Cells were stimulated with isoproterenol in the presence of the indicated concentrations of GABA. Mean ± S.E.M., n=3. Data for the “Control” condition are the same as for the “Control” presented in Figure 2. *Right,* A representative immunoblotting result confirms the expression of the Gαi-GTP bONE-GO sensor and that it does not affect expression of endogenous G proteins; multi-plasmid condition is Gαi-GTP bystander BRET sensor expressed under the same conditions as in Figure 2.

### Gαi bystander BRET sensor detects responses triggered by endogenous opioid receptors

While evidence presented above demonstrates the suitability of the bystander BRET biosensor for detecting endogenous Gαi-GTP in cells, experiments were carried out with exogenously expressed GPCRs. To test if this biosensor system was adequate for detecting G_i_ activation by endogenous GPCRs, we turned to SH-SY5Y cells, a neuroblastoma cell line that expresses endogenously the opioid receptors MOR and DOR (*32, 43*). At the same time, we set out to showcase the versatility of the biosensor by deploying it in three different formats: (1) transient transfection of the multi-plasmid design (**Fig. 5A**), (2) short-term lentiviral transduction of the bONE-GO design (**Fig. 5B**), and (3) stable expression of the bONE-GO construct (**Fig. 5C**). The purpose of this three-pronged approach was to provide other investigators with a framework of options to implement the biosensor depending on their technical resources, expertise, and preferences. Approach (1) was carried out with inexpensive transfection reagents (i.e., PEI), although it required a larger amount of cells to obtain reliable luminescence signals compared to the other approaches (see *Experimental Procedures*). For approach (2), Gαi bONE-GO-bearing lentiviral particles produced in the supernatant of HEK293T cells were applied to SH-SY5Y cells the day before BRET measurements. For approach (3), SH-SY5Y cells were transduced with lentiviral supernatants, expanded, and sensor-positive cells were then isolated by fluorescence activated cell sorting (FACS). In all three cases, BRET responses were detected upon stimulation of opioid receptors (OR) with the MOR-specific agonist DAMGO or the DOR-specific agonist SNC80 (**Fig. 5**), which were rapidly reverted upon addition of the opioid antagonist naloxone. Controls with pertussis toxin confirmed that the responses were dependent on GPCR-mediated G_i_ activation (**Fig. 5**). Taken together, these experiments indicate that the bystander BRET sensor is suitable for detecting the activation of endogenous G proteins by endogenous GPCRs when implemented in a variety of experimental formats.

**Figure 5.**
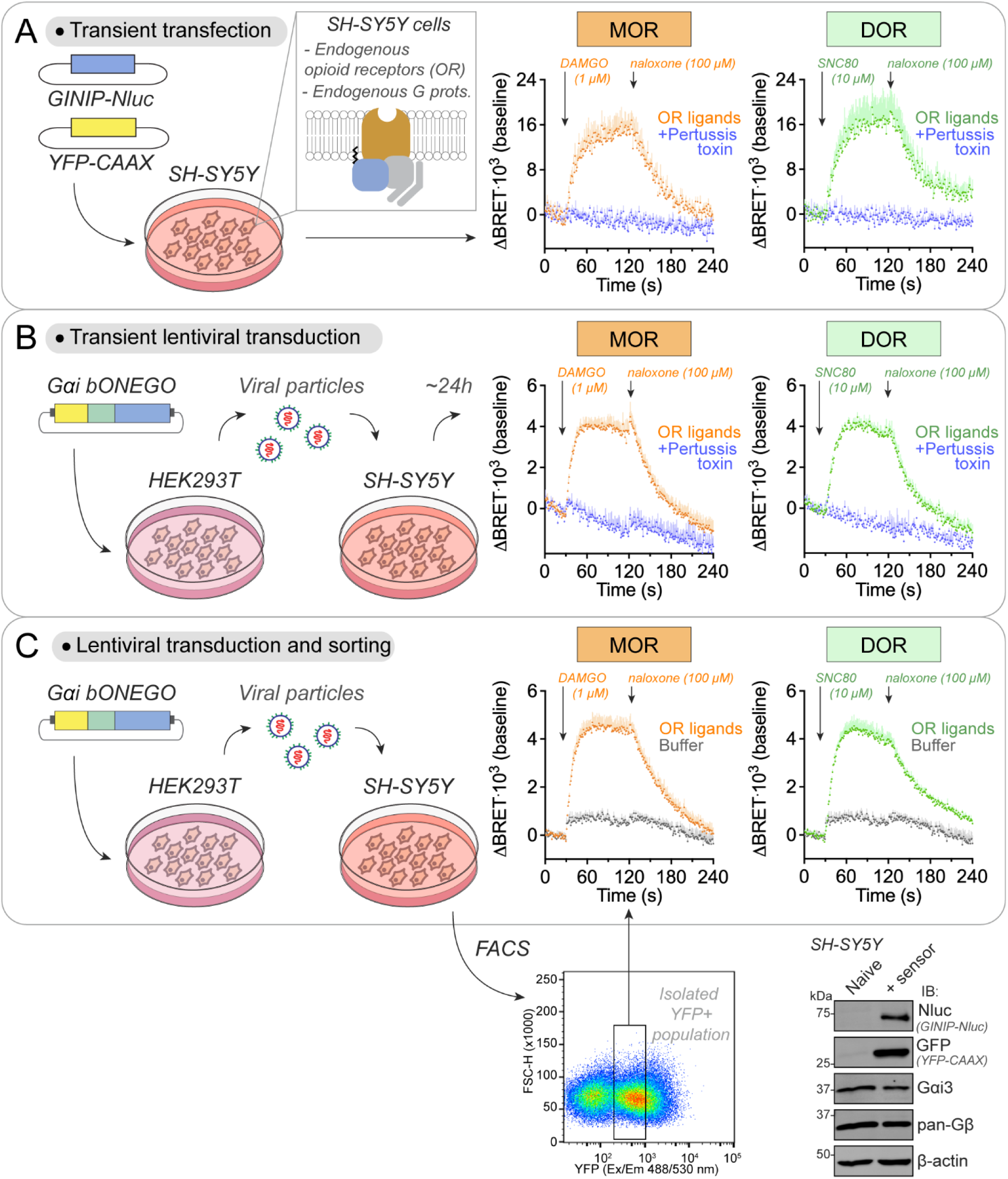
Detection of endogenous Gi activation by endogenous GPCRs in SH-SY5Y cells using a Gαi-GTP bystander BRET sensor. **(A)** Detection of endogenous Gαi activation by endogenous μ-opioid receptors (MOR) and δ-opioid receptors (DOR) in SH-SY5Y cells upon transfection with BRET sensor components. Kinetic BRET measurements were carried out in SH-SY5Y cells expressing GINIP-Nluc, and YFP-CAAX (but no exogenous G protein or GPCR) in the absence (orange for MOR, green for DOR) or presence (blue) of Pertussis toxin via PTX-S1 expression. Cells were treated with the indicated opioid receptor (OR) ligands. Mean ± S.E.M., n=6 (for MOR) or n=5 (for DOR). **(B)** Detection of endogenous Gαi activation by endogenous MOR and DOR in SH-SY5Y cells upon transient lentiviral transduction with the single-vector Gαi bONE-GO sensor construct. Kinetic BRET measurements were carried out as in (A). Mean ± S.E.M., n=5 (for MOR “OR ligands”), n=4 (for DOR “OR ligands”), n=2 (for MOR or DOR, “+ Pertussis toxin”). **(C)** Detection of endogenous Gαi activation by endogenous MOR and DOR in SH-SY5Y cells stably expressing the Gαi bONE-GO BRET sensor. SH-SY5Y cells stably expressing the Gαi bONE-GO sensor were isolated by FACS after lentiviral transduction. Kinetic BRET measurements were carried out as in (A), except that control traces (gray) were treated with buffer instead of OR ligands. Mean ± S.E.M., n=4 (for MOR), n=4 (for DOR). A representative immunoblotting result confirms expression of sensor components compared to naïve SH-SY5Y cells.

### Agonist efficacy of opioid neuropeptides on endogenous opioid receptors in SH-SY5Y cells

Next, we set out to determine the agonist efficacy of opioid neuropeptides that serve as physiological receptor ligands when detecting endogenous G protein activation in SH-SY5Y cells expressing endogenous opioid receptors. While the agonist efficacy of opioid neuropeptides like Dynorphin A, Leu-Enkephalin, Met-Enkephalin, Endomorphin-1, Endomorphin-2, and β-endorphin has been determined and annotated in the IUPHAR database (*44*), the approaches used entailed the overexpression of exogenous receptors and/or indirect readouts of activity subject to amplification (e.g., second messenger quantification). We reasoned that direct detection of G protein activity with an endogenous complement of receptors and G proteins might provide a better representation of the properties of these natural ligands under physiological conditions. For this, we stimulated SH-SY5Y cells stably expressing the Gαi bONE-GO sensor with concentrations of Dynorphin A, Leu-Enkephalin, Met-Enkephalin, Endomorphin-1, Endomorphin-2, and β-endorphin expected to saturate their cognate receptors based on their respective affinities (*44*). The six opioid neuropeptides triggered BRET responses that were comparable in magnitude to those observed upon stimulation with the synthetic MOR-specific agonist DAMGO or the synthetic DOR-specific agonist SNC80 (**Fig. 6, Fig. S2**). Since many of the opioid neuropeptides used are known to stimulate more than one opioid receptor (*45*), we envisioned an approach to isolate the response components associated to individual opioid receptors, as well as to determine their efficacy relative to an internal reference benchmark. The workflow implemented for this purpose is shown in **Fig. 6** with one representative neuropeptide (Dynorphin A), whereas the full dataset with all the opioid neuropeptides tested is presented in **Fig. S2**. The approach relied on using CTOP and ICI174,864, which are antagonists specific for the MOR and the DOR, respectively, to determine what fraction of the responses observed was mediated by each one of the receptors. Simultaneous treatment with both antagonists ablated the responses to any of the six neuropeptides, DAMGO, or SNC80, indicating that MOR and DOR, collectively, account for the responses detected in these cells (see graphs on the left in **Fig. 6** and **Fig. S2**). To isolate the MOR-specific component of the response triggered by each neuropeptide, we subtracted the response observed in the presence of the MOR-specific antagonist CTOP from the response observed under control conditions, whereas to isolate the DOR-specific component, we subtracted the response observed in the presence of the DOR-specific antagonist ICI174,864 (**Fig. 6A-B, Fig. S2A-B**). To determine the relative efficacy of each one of the opioid neuropeptides on each receptor, we compared MOR and DOR response components to those obtained upon stimulation with the full agonists DAMGO and SNC80 as internal benchmarks (**Fig. 6D, Fig. S2**). We found that most of the active opioid neuropeptides were partial agonists for the MOR and DOR (**Fig. 6D**), whereas all of them are annotated as full agonists in the IUPHAR database (*44*) (**Fig. 6E**), with the exceptions of the partial agonist annotation of Leu-Enkephalin on MOR and the lack of annotation for Endomorphin-2 on DOR (suggestive of lack of reported activity) (**Fig. 6E**). It is worth noting that Endomorphin-1 is annotated as a full agonist for DOR in the IUPHAR database (*44*), but the source reference for this annotation (*46*) does not support this claim. This suggests that Endomorphin-1 is not a DOR agonist, which is in agreement with our results showing that Endomorphin-1 lacks agonist activity on the endogenous DOR in SH-SY5Y cells (**Fig. 6D, Fig S2C**). These results obtained using the Gαi bONE-GO sensor in SH-SY5Y cells also contrast with some evidence using other BRET-based biosensors that detect G protein activity directly, like TRUPATH or ONE-GO, which also indicated full agonist activity of these opioid neuropeptides on the MOR exogenously expressed in HEK293 cells (*30, 33*). To more rigorously characterize this difference with the endogenous responses observed in SH-SY5Y cells, we measured G protein activation with the previously described Gαi1 ONE-GO sensor (*33*) in HEK293T cells expressing either exogenous MOR or DOR upon stimulation with saturating concentrations of the opioid neuropeptides. We found that all the opioid neuropeptides that elicited a response did so as full agonists, as assessed by direct comparison with the MOR- or DOR-specific full agonists DAMGO or SNC80, respectively (**Fig. 6F**). Overall, these results indicate that the pharmacological properties of natural opioid neuropeptides can be distorted when the signaling components of the system are not expressed at endogenous levels, and that the Gαi bONE-GO sensor might provide a better representation of physiological signaling responses.

**Figure 6.**
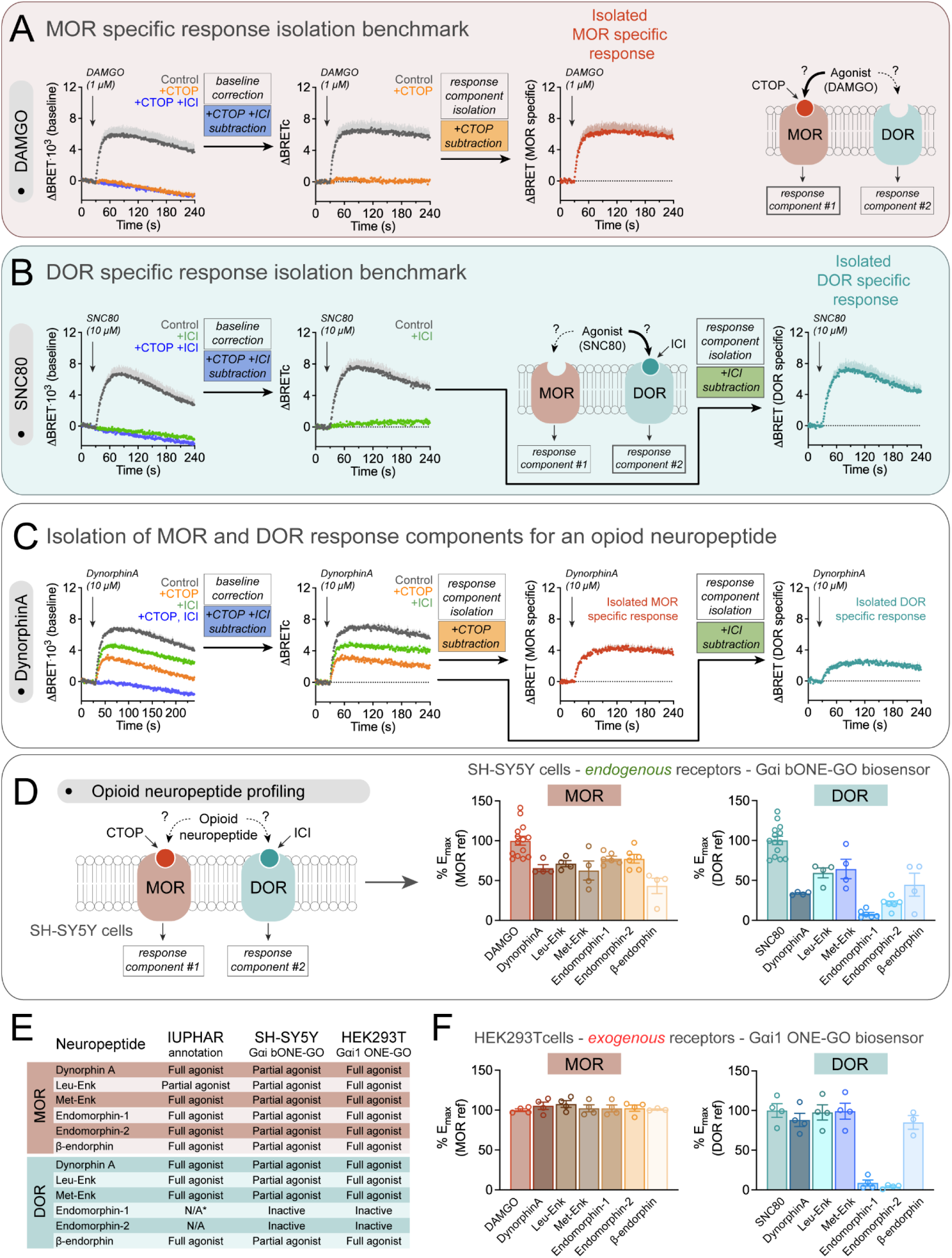
bONE-GO reveals partial agonism of opioid neuropeptides on endogenous receptors. **(A)** Benchmarking of full agonist MOR-specific response in SH-SY5Y cells with the Gαi bONE-GO sensor. Kinetic BRET measurements were carried out in SH-SY5Y cells stably expressing the Gαi bONE-GO sensor in the absence (Control, grey) or presence of 10 μM CTOP (+CTOP, orange), or 10 μM CTOP and 100 μM ICI174,864 (+CTOP +ICI, blue), followed by stimulation with DAMGO. To isolate the MOR-specific response component, first the baseline trace obtained in the presence of CTOP and ICI (+CTOP +ICI) was subtracted from other measurements, followed by the subtraction of the +CTOP trace from the control. Mean ± S.E.M., n=4. **(B)** Benchmarking of full agonist DOR-specific response in SH-SY5Y cells with the Gαi bONE-GO sensor. Kinetic BRET measurements were carried out as in (A) in the absence (Control, grey) or presence of 100 μM ICI174,864 (+ICI, green) or 10 μM CTOP and 100 μM ICI174,864 (+CTOP +ICI, blue) following stimulation with SNC80. To isolate the DOR-specific response component, first the baseline trace obtained in the presence of CTOP and ICI (+CTOP +ICI) was subtracted from other measurements, followed by the subtraction of the +ICI trace from the control. Mean ± S.E.M., n=4. **(C)** Isolation of MOR- and DOR-specific responses elicited by Dynorphin A in SH-SY5Y cells. Kinetic BRET measurements were carried out as in (A) in the absence (Control, grey) or presence of 10 μM CTOP (+CTOP, orange), 100 μM ICI174,864 (+ICI, green), or 10 μM CTOP and 100 μM ICI174,864 (+CTOP +ICI, blue) followed by stimulation with Dynorphin A. To isolate the MOR- and DOR-specific response components, data were processed as in (A) for the MOR-specific component or as in (B) for the DOR-specific component. Mean ± S.E.M., n=4. **(D)** Assessment of agonist efficacy of opioid neuropeptides on endogenous opioid receptors in SH-SY5Y cells using Gαi bONE-GO. *Left,* diagram representation of opioid neuropeptide profiling for MOR- or DOR-specific response components. *Right*, the isolated MOR- and DOR-specific response components of each opioid neuropeptide tested in this figure and **Fig. S2** at the indicated concentrations were expressed relative to the maximal responses (%E_max_) elicited by DAMGO or SNC80 for MOR and DOR, respectively. Mean ± S.E.M., n=3-14. **(E)** Table summarizing agonist efficacy of opioid neuropeptides for ORs based on IUPHAR annotation or detection with Gαi bONE-GO (from panel **D**) or Gαi1 ONE-GO (from panel **F**). N/A; no annotation, presumably inactive. *Although Endomorphin-1 is annotated as a full agonist for DOR in the IUPHAR database, the evidence in the reference provided in the database indicates that it is inactive. **(F)** Assessment of agonist efficacy of opioid neuropeptides on exogenous opioid receptors in HEK293T cells using Gαi1 ONE-GO. Endpoint BRET experiments were carried out in HEK293T cells expressing Gαi1 ONE-GO and either MOR or DOR, as indicated, following stimulation with 1 μM DAMGO, 10 μM SNC80, 10 μM Dynorphin A, 10 μM Leu-Enkephalin, 10 μM Met-Enkephalin, 10 μM Endormorphin-1, 10 μM Endormorphin-2, or 10 μM β-endorphin. Responses were expressed relative to the maximal responses (%E_max_) elicited by DAMGO or SNC80 for MOR and DOR, respectively. Mean ± S.E.M., n=3-4. Panels A and B, and C contain data also presented in **Figure S2.**

### Gαi bONE-GO sensor reports activation of adenosine receptors in mouse glial cells

While detection of responses triggered by endogenous receptors, as shown above with the Gαi bONE- GO sensor for opioid receptors in SH-SY5Y cells, is a desirable feature towards dissecting physiologically-relevant signaling mechanisms, cell lines do not always recapitulate the behavior and characteristics of non-immortalized cells. Thus, we set out to assess if the Gαi bONE-GO sensor could be successfully implemented in primary cells. For this, we transduced mouse cortical astroglial cells with a lentivirus for the expression of the Gαi bONE-GO sensor and stimulated them with adenosine, which is known to stimulate A1 purinergic receptors in these cells (*33, 47*) (**Fig. 7**). We found that adenosine stimulation led to robust and concentration-dependent responses (**Fig. 7**). Adenosine responses were completely suppressed after treatment of the cells with pertussis toxin, and not recapitulated in cells expressing a Gαi bONE-GO construct bearing a mutation in GINIP (W139A) that disrupts G protein binding (**Fig. 7**), confirming the expected specificity of the BRET response observed with the Gαi bONE-GO sensor. These results indicate that the Gαi bONE-GO sensor is suitable for the characterization of responses elicited by endogenous GPCRs and endogenous G proteins in primary cells.

**Figure 7.**
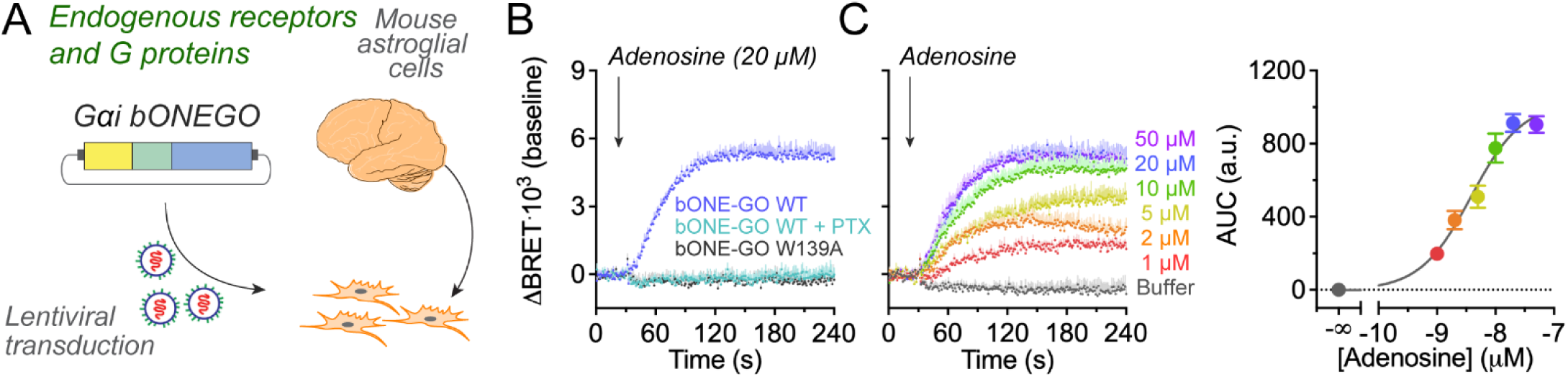
Detection of endogenous responses to adenosine in astroglial cells using Gαi bONE-GO sensor. **(A)** Diagram depicting lentiviral transduction of cultured primary mouse astroglial cells. **(B)** Detection of endogenous Gαi activation by endogenous adenosine receptors using Gαi bONE-GO. Kinetic BRET measurements were carried out in primary mouse astroglial cells upon lentiviral transduction with Gαi bONE-GO WT (blue, cyan) or Gαi bONE-GO W139A (grey) in the absence (blue, grey) or presence (cyan) of overnight treatment with Pertussis toxin (PTX). Cells were treated with adenosine as indicated. Mean ± S.E.M., n=3. **(C)** Gαi bONE-GO detects concentration-dependent activation of endogenous Gαi by endogenous adenosine receptors. Kinetic BRET measurements were carried out with Gαi bONE-GO WT as in (B). Cells were treated with the indicated concentrations of adenosine or buffer. The area under curve (AUC) of the responses detected in the kinetic traces was calculated for each concentration of adenosine, and plotted as a semi-log graph on the right. Mean ± S.E.M., n=3.

## DISCUSSION

The main advance provided by this work is the development of a biosensor design, Gαi bONE-GO, that allows for direct measurement of endogenous Gαi-GTP generated upon stimulation of endogenous GPCRs and the demonstration of its versatile implementation across experimental systems to reveal more physiologically-relevant information on G protein signaling. This sensor design improves the dynamic range over what was observed with previously developed Gαi*-BERKY biosensors also capable of measuring activity with endogenous GPCRs and G proteins, while lacking interference with signaling to downstream effectors and allowing for deployment in different assay formats and across different cell types, including primary cell cultures. Thus, this design also overcomes some limitations of other biosensor platforms like ONE-GO or EMTA, which may compromise G protein function and/or are not suitable for implementation in primary cells. The significance of having an approach to faithfully investigate endogenous Gαi activation in response to endogenous GPCR stimulation was showcased by revealing that several natural opioid neuropeptides work as partial agonists under endogenous expression conditions, contrary to observations obtained by direct comparison with another biosensor, Gαi1 ONE-GO, using overexpressed receptors and exogenous G proteins, which revealed full agonist activity. Overall, the Gαi bONE-GO biosensor represents a design that combines the desirable features of previously developed G protein sensor platforms while overcoming their limitations.

There are three key features of the Gαi bONE-GO design that are critical for its improved performance: (1) using GINIP as the detector module; (2) leveraging the principle of bystander BRET; and (3) assembling all biosensor components into a single vector. Using GINIP as the detector module not only increases sensitivity and dynamic range because of its high affinity for Gαi-GTP, but also contributes to the lack of interference with downstream signaling because it does not directly alter nucleotide binding or hydrolysis by the G protein (*39*). As for leveraging the principle of bystander BRET, one advantage is that it allows for detection of Gαi-GTP without the need to fuse the G protein to bulky tags or to express it as an exogenous protein. It is also possible that the use of bystander BRET as readout is conducive to a better dynamic range of detection, since the acceptor-to-donor ratio at the plasma membrane might be large. Finally, the assembly of all biosensor components into a single vector akin to the recently described ONE-GO sensor design (*33*) allows for a reduction in the overall level of expression of GINIP, thereby further mitigating interference with downstream signaling, and facilitates implementation in different experimental systems, even in difficult to transfect cell types, by virtue of requiring the delivery of a single genetic payload. Overall, in developing the Gαi bONE-GO design we overcame limitations of existing G protein activity biosensors by leveraging a combination of their desirable features with a better detector module, an approach that may serve as a template for the future development of analogous biosensors for other G protein subtypes.

Implementing Gαi bONE-GO to detect endogenous G protein activation by endogenously expressed GPCRs holds the promise of revealing new insights into how this signaling mechanism occurs under native conditions, as illustrated by our results profiling the action of opioid neuropeptides. Our results with endogenous receptors and G proteins expressed in SH-SY5Y cells using the Gαi bONE-GO sensor revealed that most of them act as partial agonists instead of displaying the full agonism annotated in the IUPHAR database (*44*). One potential explanation for this discrepancy is that IUPHAR database annotations rely largely on assays that measure amplified second messenger responses. However, it is also likely that receptor overexpression is a major contributor to the observed differences, since full agonism is also detected when using biosensors that directly measure G protein activity like TRUPATH (*30*) or ONE-GO (*33*) in HEK293 cells overexpressing opioid receptors. It is therefore conceivable that either the amplification associated with the measurement of second messengers and/or the excess of receptor skews the responses observed compared to the direct measurement of G protein activity in a system with native receptor-G protein stoichiometry. These observations also resonate with recent findings supporting that context-dependence is a prevalent feature of G protein activation by endogenous GPCRs (*33*). Our findings also reinforce the cautionary message of a recent report showing that biosensor responses for GPCRs coupled to G_i/o_ proteins are not only different between cell lines and primary neurons, but are also influenced by the type of exogenous G protein subunits required to assemble the biosensor system (*31*). Overall, the context-dependence of GPCR responses, like that shown here for opioid receptors, impacts the translatability of pharmacological profiling results *in vitro* into the expected effects of a given drug *in vivo* (*48, 49*). More specifically, our findings have important implications in the context of the development of novel opioid analgesics with diminished side effects, an area that remains controversial (*50*). While some evidence suggests that preferential activation of G proteins over arrestins (i.e., G protein-bias) by MOR has improved safety profiles (*18, 51*), others have put this into question (*52–54*). Interestingly, one report has provided evidence that the reduced side effects of several G protein-biased opioid agonists can be explained by their low intrinsic efficacy (*53*). Given that our results reveal that efficacy is a function of the system and/or experimental conditions, it will be important in the future to critically assess the action of existing or new opioid analogs under physiologically relevant conditions. Approaches like the one developed here hold the promise of enabling this type of assessment.

An attractive feature of the Gαi bONE-GO sensor design is its versatility in terms of implementation, as showcased by the variety of systems and formats described in this work. In addition to making it easy to scale up throughput in experiments in HEK293T cells by easily transfecting a single plasmid, viral transduction of a single payload makes it feasible to use the Gαi bONE-GO sensor in cell types that are not efficiently transfected, as exemplified here with SH-SY5Y cells and astroglial cells. In the context of drug discovery, this could increase the success of translating pharmacological properties *in vitro* to desired outcomes *in vivo* by establishing an intermediary step of testing the compounds under development in primary cells relevant to the particular indication. For example, one could use the same readout (i.e., Gαi bONE-GO sensor) to directly assess whether the responses observed in a relevant cell type expressing endogenously the receptor of interest resemble those obtained in a cell line expressing the receptor exogenously. The option of making stable cell lines to monitor endogenous GPCR responses, as we illustrated here with SH-SY5Y cells, could also be attractive for high-throughput drug screening campaigns, in which the variability associated with transient transfections is detrimental. Another aspect related to the versatility of the Gαi bONE-GO sensor is that its design allows for relatively easy customization. For example, the bystander BRET acceptor module could be targeted to different subcellular locations, like endosomes or the Golgi apparatus, by replacing the polybasic-CAAX sequence with targeting sequences suitable for the alternative locations of interest (*55, 56*). This could be useful to directly dissect the spatiotemporal pattern of Gαi activation, an area of current interest for GPCRs in general and for G_i_-coupled opioid receptors in particular. Opioid receptors can be activated in different subcellular locations and timescales depending on the nature of the ligand, and signaling from each location might lead to different functional outcomes (*57–60*). Future iterations of the bONE-GO design may be of use for capturing the formation of active G proteins in different subcellular compartments by taking advantage of the bystander design of the sensor.

In summary, Gαi bONE-GO combines the desirable design features of other existing biosensor platforms, while overcoming some of their limitations, to provide high fidelity detection of endogenous GPCR-G protein signaling with the flexibility for use in a wide variety on contexts. By providing proof-of-principle evidence for its implementation in diverse experimental formats and for the conceptual advances that can be obtained through it, we hope to entice other investigators to leverage this system in order to advance in the field of GPCR signaling.

## EXPERIMANTAL PROCEDURES

### Plasmids

The plasmids for the expression of GINIP-Nluc in mammalian cells via transfection (p3xFLAG-CMV-14-GINIP-Nluc) have been described previously (*39*). The plasmid encoding YFP-CAAX, consisting of Venus followed by the last 25 amino acids of human KRas4b including the polybasic regions and CAAX prenylation box, was a gift from Nevin Lambert (*55*). The plasmid for mammalian expression of the long isoform of the human Dopamine 2 receptor (pcDNA3.1(+)-FLAG-D2DR) was provided by A. Kovoor (University of Rhode Island) (*61*). The plasmid encoding rat α2_A_-AR (pcDNA3-α2_A_-AR) was provided by Joe Blumer (Medical University of South Carolina) has been described previously (*62*). The plasmids encoding rat GABA_B_R subunits (pcDNA3.1(+)-GABA_B_R1a and pcDNA3.1(+)-GABA_B_R2) were a gift from Paul Slessinger (Ichan School of Medicine Mount Sinai, NY). The plasmid encoding mouse MOR (pcDNA3.1-MOR-FLAG) has been described previously (*28*). The plasmids encoding β2AR (cat#14697; (*63*)), PAR1 (cat#53226; (*64*)), XE100 Pertussis Toxin A promoter (called PTX-S1 where applicable; cat#16678), were obtained from Addgene, as well as the plasmids psPAX2 (cat#12259), pMD2.G (cat#12259) used for lentiviral packaging. The plasmids encoding human DOR (cat#OPRD100000) or M3R (cat#MAR030TN00) were obtained from the cDNA Resource Center at Bloomsburg University. The plasmid for expression of the Gαi*-BERKY3 biosensor (pcDNA3.1-Gαi*-BERKY3) was generated in a previous study (*65*). Plasmids encoding Gαs ONE-GO (pLVX-CMV-Gαs-99V-IRES*-KB1691-Nluc-T2A-Ric-8B), Gαq ONE-GO (pLVX-CMV-Gαq-V-IRES*-GRK2^RH^-Nluc), Gα13 ONE-GO (pLVX-CMV-Gα13-V-IRES*-PRG^RH^-Nluc), and Gαi1 ONE-GO (pLVX-CMV-Gαi1-V-IRES*-KB1753-Nluc) were described previously (*33*). The plasmid encoding Glosensor 22F was acquired from Promega (cat#E2301). The plasmid for the expression of the Gαi bONE-GO biosensor (pLVX-CMV-YFP-CAAX-IRES*-GINIP-Nluc) was generated by replacing the IRES-Hyg cassette between the BamHI and MluI sites of pLVX-IRES-Hyg with YFP-CAAX, IRES*, and GINIP-Nluc using Gibson assembly. The sequences encoding the YFP-CAAX and GINIP-Nluc cassettes were amplified from plasmids described above, and IRES* is a previously described sequence that leads to lower expression of the gene of interest downstream of it relative to the gene of interest right downstream of the CMV promoter (*33, 66*). All point mutations were generated using QuikChange II following the manufacturer’s instructions (Agilent, Cat#200523).

### Bioluminescence Resonance Energy Transfer (BRET) measurements in HEK293T cells

HEK293T cells (ATCC, cat#CRL-3216) were grown at 37°C, 5% CO_2_ in DMEM (Gibco, cat#11965-092) supplemented with 10% FCS (Hyclone, cat#SH30072.03), 100 units/ml penicillin, 100 μg/ml streptomycin, and 2 mM L-glutamine (Corning, cat#30-009-CI).

Approximately 400,000 HEK293T cells were seeded on each well of 6-well plates coated with 0.1% (w/v) gelatin, and transfected ∼24 hr later using the calcium phosphate method. Cells were transfected, in the combinations indicated in the figures, with plasmids encoding the following constructs (DNA amounts in parentheses): GABA_B_R1a (0.2 μg), GABA_B_R2 (0.2 μg), α2_A_-AR (0.2 μg), D2R (0.2 μg), MOR (0.2 μg), β2AR (0.2 μg), M3R (0.02 μg), PAR1 (0.2 μg), DOR (0.2 μg), YFP-CAAX (1 μg), GINIP-Nluc (0.05 μg), PTX-S1 (0.2 μg), Gαs ONE-GO (0.08 μg), Gαq ONE-GO (0.05 μg), Gα13 ONE-GO (0.05 μg), Gαi*-BERKY3 (0.01 μg), Gαi1 ONE-GO (0.05 μg), and Gαi bONE-GO (0.025 μg). Total DNA amount per well was equalized by supplementing with empty pcDNA3.1 as needed. Cell medium was changed 6 hr after transfection.

For kinetic BRET measurements, approximately 18-22 hr after transfection, cells were washed and gently scraped in room temperature Phosphate Buffered Saline (PBS; 137 mM NaCl, 2.7 mM KCl, 8 mM Na_2_HPO_4_, and 2 mM KH_2_PO_4_), centrifuged (5 minutes at 550 × *g*), and resuspended in 750 μl of BRET buffer (140 mM NaCl, 5 mM KCl, 1 mM MgCl_2_, 1 mM CaCl_2_, 0.37 mM NaH_2_PO_4_, 24 mM NaHCO_3_, 10 mM HEPES, and 0.1% glucose, pH 7.4). Approximately 25-50 μl of cells were added to a white opaque 96-well plate (Opti-Plate, PerkinElmer Life Sciences, cat#6005290). BRET buffer was added to a final volume of 100 μl and then mixed with the nanoluciferase substrate Nano-Glo (Promega, cat#N1120, final dilution 1:200) before measuring luminescence. Luminescence signals at 450 ± 40 and 535 ± 15 nm were measured at 28°C every 0.96 s in a BMG Labtech POLARStar Omega plate reader. Agonists were added as indicated in the figures during the recordings using built-in injectors. BRET was calculated as the ratio between the emission intensity at 535 nm divided by the emission intensity at 450 nm, followed by multiplication by 10^3^. Kinetic traces are represented as change in BRET after subtraction of the baseline signal measured for 30 s before GPCR stimulation [ΔBRETꞏ10^3^ (baseline)].

For endpoint BRET measurements to determine concentration dependence curves, cells were scraped and resuspended in BRET buffer as described above except that they were resuspended in 300 μl BRET buffer. Twenty μl of GABA, brimonidine, dopamine, DAMGO, SNC80, Dynorphin A, Leu-Enkephalin, Met-Enkephalin, Endomorphin-1, Endomorphin-2, or β-endorphin diluted in BRET buffer at 5X the final concentration desired in the assay were added to wells of a white opaque 96-well plate and further diluted with 35 μl of BRET buffer. Next, 22.4 μl of BRET buffer containing the luciferase substrate CTZ400a (GoldBio, cat#C-320-1; 10 μM final concentration) was added to wells. Cell stimulation was initiated by adding 22.4 μl of cell suspension to wells containing the agonists and the luciferase substrate. Luminescence signals at 450 ± 40 and 535 ± 15 nm were measured at 28°C every minute for 5 minutes in a BMG Labtech POLARStar Omega plate reader with a signal integration time of 0.32 s for each measurement. BRET was calculated as the ratio between the emission intensity at 535 nm divided by the emission intensity at 450 nm for each time point, and the two values obtained at 4 and 5 minutes were averaged and multiplied by 10^3^. BRET data are presented as the change in BRET relative to a condition without agonist [ΔBRETꞏ10^3^ (no agonist)]. In some cases, the final values were fit to a curve using a 3-parameter sigmoidal curve-fit in Prism (GraphPad).

### Luminescence-based cAMP measurements in HEK293T cells

Culture conditions for HEK293T cells are described above in *‘Bioluminescence Resonance Energy Transfer (BRET) measurements in HEK293T cells*.*’*

Approximately 300,000 HEK293T cells were seeded on each well of 6-well plates coated with 0.1% (w/v) gelatin, and transfected ∼24 hr later with plasmids using the calcium phosphate method. Cells were transfected, in the combinations indicated in the figures, with plasmids encoding the following constructs (DNA amounts in parentheses): GABA_B_R1a (0.2 μg), GABA_B_R2 (0.2 μg), Glosensor 22F (0.8 μg), YFP-CAAX (1 μg), GINIP-Nluc WT (0.05 μg), and Gαi bONE-GO (0.025 μg), supplemented with pcDNA3.1 to equalize total amount of DNA per well and reach a minimum of 2 μg of total transfected DNA for all experiments. Cell medium was changed 6 hr after transfection.

For kinetic measurements, approximately 18-22 hr after transfection, cells were washed and gently scraped in room temperature PBS, centrifuged (5 minutes at 550 × *g*), and resuspended in 750 μl Tyrode’s solution (140 mM NaCl, 5 mM KCl, 1 mM MgCl_2_, 1 mM CaCl_2_, 0.37 mM NaH_2_PO_4_, 24 mM NaHCO_3_, 10 mM HEPES and 0.1% glucose, pH 7.4). Two-hundred μl of cells were mixed with 200 μl of 5 mM D-luciferin K^+^ salt (GoldBio, cat#LUCK-100) diluted in Tyrode’s solution and incubated at 28°C for 15 minutes. Ninety μl of cells pre-incubated with D-luciferin were added to a white opaque 96-well plate before measuring luminescence without filters at 28°C every 10 s in a BMG Labtech POLARStar Omega plate reader. Agonists were added as indicated in the figures during the recordings using built-in injectors. Kinetic traces are represented as the percentage of the maximum response after stimulation with isoproterenol only [cAMP (% isoproterenol max)].

For concentration-response curves, cells were washed and scraped as above, except that they were resuspended in 300 μl Tyrode’s solution. Two-hundred and forty μl of cells were mixed with 240 μl of 5 mM D-luciferin K+ salt diluted in Tyrode’s solution and incubated at 28°C for 15 minutes. Twenty μl of different amounts of GABA diluted in Tyrode’s solution at 4X the final concentration desired in the assay were added to wells of a white opaque 96-well plate, and further diluted by addition of 37.6 μl of Tyrode’s solution. GABA stimulations were initiated at room temperature by addition of 22.4 μl of the cell suspension pre-incubated with D-luciferin to the wells, and 2 minutes later 20 μl of 500 nM isoproterenol (100 nM final concentration) diluted in Tyrode’s solution were added. Immediately following addition of isoproterenol, luminescence measurements without filters were taken at 28°C for 19 minutes in 30 s intervals using a BMG Labtech POLARStar Omega plate reader with a signal integration time of 0.20 s for each measurement. For each concentration of GABA, response values were calculated by averaging the 3 time points around the peak of the kinetic trace (270, 300, and 330 s after start of measurement) and normalizing them as a percentage of the maximum response in the absence of GABA [cAMP (% isoproterenol max)]. Where indicated, the IC_50_ values and concentration dependence curves were determined by using a 3-parameter sigmoidal curve-fit in Prism (GraphPad).

At the end of experiments, a separate aliquot of the same pool of cells used for the measurements was centrifuged for 1 minute at 14,000 × *g*, and pellets stored at −20°C for subsequent immunoblot analysis (see “*Protein electrophoresis and Immunoblotting*” section below).

### Bioluminescence Resonance Energy Transfer (BRET) measurements in SH-SY5Y cells

SH-SY5Y cells (ATCC cat#CRL-2266) were grown at 37°C, 5% CO_2_ in DMEM supplemented with 100 U/ml penicillin, 100 μg/ml streptomycin, 2 mM L-glutamine, and 15% heat-inactivated FCS (Hyclone, cat#SH30072.03).

For experiments using transient transfection of naïve SH-SY5Y cells with the multi-plasmid Gαi-GTP bystander BRET sensor (**Fig. 5A**), approximately 800,000 SH-SY5Y cells were seeded on each well of 6-well plates coated with 0.1% (w/v) gelatin and transfected ∼24 hr later with plasmids using the polyethylenimine (PEI) method (*67*). The following plasmids were transfected using a 1:2 ratio of DNA to PEI (DNA amounts in parentheses): YFP-CAAX (1 μg), GINIP-Nluc WT (0.05 μg), and PTX-S1 (0.2 μg). Total DNA amount per well was equalized by supplementing with empty pcDNA3.1 to also reach a minimum of 2.5 μg of total transfected DNA. Cell medium was changed 6 hr after transfection, and approximately 16-24 h after transfection, cells were washed and gently scraped in room temperature PBS, centrifuged (5 minutes at 550 × *g*), and resuspended in 375 μl of BRET buffer. Fifty μl of cells were added to a white opaque 96-well plate, followed by addition of 50 μl of BRET buffer and the nanoluciferase substrate Nano-Glo (final dilution 1:200) before measuring luminescence. Luminescence signals at 450 ± 40 and 535 ± 15 nm were measured at 28 °C every 0.96 s in a BMG Labtech POLARStar Omega plate reader, and BRET was calculated as the ratio between the emission intensity at 535 nm divided by the emission intensity at 450 nm, followed by multiplication by 10^3^. Agonists were added as indicated in the figures during the recordings using built-in injectors. Kinetic traces are represented as the change in BRET after subtraction of the baseline signal measured for 30 s before GPCR stimulation [ΔBRETꞏ10^3^ (baseline)].

For experiments using transient lentiviral transduction of SH-SY5Y cells with Gαi bONE-GO BRET sensor (**Fig. 5B**), supernatants containing viral particles were first generated in HEK293T cells as described next. Approximately 400,000 HEK293T cells were seeded on each well of 6-well plates coated with 0.1% (w/v) gelatin, and transfected ∼24 hr later with plasmids encoding the following components using the PEI method at a 1:2 ratio of DNA to PEI (DNA amounts in parentheses): Gαi bONE-GO (1.8 μg), psPAX2 (1.2 μg), and pMD2.g (0.75 μg). Cell medium was changed 6 hr after transfection. Lentivirus-containing media was collected 24 hr and 48 hr after transfection, pooled, centrifuged for 5 minutes at 1500 × *g*, and filtered through a 0.45-μm surfactant-free cellulose acetate (SFCA) membrane filter (Corning, cat#431220). These supernatants (∼4 ml collected per well of cultured cells), were stored at 4°C for up to 7 days before using them to transduce SH-SY5Y cells. In parallel, approximately 800,000 SH-SY5Y cells were seeded on each well of 6-well plates coated with 0.1% (w/v) gelatin and transduced ∼24 hr later by replacing cell media with 2 ml of a 1:1 mix of lentivirus-containing supernatants and fresh complete medium supplemented with 6 µg/ml of polybrene (Tocris Bioscience, cat#7711/10) to enhance transduction efficiency. Virus-containing medium was replaced by fresh medium 6 hr later. In some conditions, the change of media was accompanied by the addition of 0.1 μg/ml pertussis toxin (List Biological Labs, cat#179A) to wells. Approximately 18-22 hr after the change to fresh medium, cells were washed and gently scraped in room temperature PBS, centrifuged (5 minutes at 550 × *g*), and resuspended in 750 μl of BRET buffer. Fifty μl of cells were added to a white opaque 96-well plate, followed by addition of 50 μl of BRET buffer and the nanoluciferase substrate Nano-Glo (final dilution 1:200) before measuring luminescence. Luminescence signals at 450 ± 40 and 535 ± 15 nm were measured at 28°C every 0.96 s in a BMG Labtech POLARStar Omega plate reader, and BRET was calculated as the ratio between the emission intensity at 535 nm divided by the emission intensity at 450 nm, followed by multiplication by 10^3^. Agonists were added as indicated in the figures during the recordings using built-in injectors. Kinetic traces are represented as change in BRET after subtraction of the baseline signal measured for 30 s before GPCR stimulation [ΔBRETꞏ10^3^ (baseline)].

### BRET measurements in SH-SY5Y cells stably expression Gαi bONE-GO

SH-SY5Y cells stably expressing Gαi bONE-GO were generated by lentiviral transduction followed by Fluorescence-Activated Cell Sorting (FACS) as described next. Approximately 800,000 SH-SY5Y cells, cultured as described above in ‘*Bioluminescence Resonance Energy Transfer (BRET) measurements in SH-SY5Y cells’,* were seeded on 35 mm tissue culture plates and transduced ∼24 hr later by replacing cell medium with 2 ml of a 1:1 mix of lentivirus-containing supernatants (collected as described above) and fresh complete medium supplemented with 6 µg/ml of polybrene. Virus-containing medium was replaced by fresh medium 48 hr later. Cells were expanded to multiple 10 cm plates as the starting material for FACS. For cell sorting, SH-SY5Y stable cells were detached by trypsin, resuspended in complete medium, and counted such that 7.5 x10^6^ cells were transferred to a 15 ml conical tube. Cells were washed 3 times with 10 ml cold PBS by cycles of centrifugation (3 minutes at 300 × *g*), aspiration, and resuspension. Cells were resuspended in 1.5 ml cold PBS and stored on ice for 3 hr while carrying out sorting protocol. A subset of the trypsinized SH-SY5Y stable cells were resuspended in complete DMEM containing DAPI (1 μg/ml), washed as described above, and used for selecting fluorescence gates. Cell sorting was performed on FACSAria II SORP (BD Bioscience), and the 488_ex_/530_em_ nm fluorescence channel (Voltage: 225 nV) was used for positive selection. Approximately 3.5 x10^5^ cells with fluorescence intensity from 200 to 1000 were collected as *“isolated YFP+ population”* (**Fig. 5C**), and seeded in a 6-well plate with complete DMEM for expansion. Culture conditions for the SH-SY5Y stable cell line were the same as described for naïve SH-SY5Y cells.

For kinetic BRET measurements using SH-SY5Y cells stably expressing Gαi bONE-GO BRET sensor (**Fig. 5C**), approximately 800,000 cells were seeded on 6 cm plates coated with 0.1% (w/v) gelatin. Approximately 18-22 hr later, cells were washed and gently scraped in room temperature PBS, centrifuged (5 minutes at 550 × *g*), and resuspended in 750 μl of BRET buffer. Fifty μl of cells were added to a white opaque 96-well plate, followed by addition of 50 μl of BRET buffer and the nanoluciferase substrate Nano-Glo (final dilution 1:200) before measuring luminescence. Luminescence signals at 450 ± 40 and 535 ± 15 nm were measured at 28 °C every 0.96 s in a BMG Labtech POLARStar Omega plate reader and BRET was calculated as the ratio between the emission intensity at 535 nm divided by the emission intensity at 450 nm, followed by multiplication by 10^3^. Agonists were added as indicated in the figures during the recordings using built-in injectors. Kinetic traces are represented as change in BRET after subtraction of the baseline signal measured for 30 s before GPCR stimulation [ΔBRETꞏ10^3^ (baseline)], except for some experiments described next.

Calculation of the pharmacologically isolated MOR- and DOR-specific components for opioid neuropeptide responses (**Fig. 6, Fig. S2**) was performed as follows. First, the trace obtained in the presence of both CTOP and ICI174,864 (ICI) was subtracted from the other conditions tested (Control, CTOP only, or ICI only) to obtain a baseline correction. Next, to isolate the MOR-specific response component, the trace obtained for each agonist condition in the presence of CTOP was subtracted from the Control trace (no inhibitors). Similarly, for the DOR-specific response component, the trace obtained for each agonist condition in the presence of ICI was subtracted from the Control trace. To obtain the data presented in **Fig. 6D**, each of the corrected and isolated OR-specific responses was quantified as the area under curve (AUC), and normalized to a maximal response (%E_max_) obtained with a full agonist for either MOR (DAMGO) or DOR (SNC80). To calculate the AUC, the total area was calculated in Prism (GraphPad) between the isolated MOR- or DOR-specific response components and y=0.

At the end of some experiments, a separate aliquot of the same pool of cells used for the measurements was centrifuged for 1 minute at 14,000 × *g* and pellets stored at −20°C for subsequent immunoblot analysis (see “*Protein electrophoresis and Immunoblotting*” section below).

### Protein electrophoresis and immunoblotting

Pellets of HEK293T or SH-SY5Y stable cells were resuspended with cold lysis buffer (20 mM HEPES, 5 mM Mg(CH_3_COO)_2_, 125 mM K(CH_3_COO), 0.4% (v:v) Triton X-100, 1 mM DTT, 10 mM β-glycerophosphate, 0.5 mM Na_3_VO_4_, supplemented with a protease inhibitor cocktail [Sigma, cat#S8830], pH 7.4) and incubated on ice for 10 minutes with intermittent vortexing. Lysates were cleared by centrifugation (10 minutes at 14,000 × *g*, 4°C) and quantified by Bradford (Bio-Rad, cat#5000205). Samples were then boiled for 5 minutes in Laemmli sample buffer. Proteins were separated by SDS-PAGE and transferred to PVDF membranes, which were blocked with 5% (w/v) nonfat dry milk in Tris Buffered Saline (TBS; 20 mM Tris-HCl and 150 mM NaCl), followed by incubation with primary antibodies diluted in 2.5% (w/v) nonfat dry milk in TBS supplemented with 0.1% (w/v) Tween-20 (TBS-T) and 0.05% (w/v) sodium azide. Secondary antibodies were diluted in 2.5% (w/v) nonfat dry milk in TBS-T. The primary antibodies used were the following (species, source, and dilution factor indicated in parenthesis): GFP (mouse, Clontech cat# 632380, 1:2,000); Gαi3 (rabbit, Aviva Cat#OAAB19207, 1:1,000); Gβ (mouse, Santa Cruz Biotechnology cat# sc-166123; 1:250); β-actin (rabbit, LI-COR Cat#926-42212; 1:1,000); Nluc (mouse, Promega cat# N700A; 1:500). The following secondary antibodies were used at a 1:10,000 dilution (species and vendor indicated in parenthesis): anti-mouse Alexa Fluor 680 (goat, Invitrogen cat# A21058); anti-mouse IRDye 800 (goat, LI-COR cat# 926-32210); anti-rabbit DyLight 800 (goat, Thermo cat# 35571). Infrared imaging of immunoblots was performed according to manufacturer’s recommendations using an Odyssey CLx infrared imaging system (LI-COR Biosciences). Images were processed using Image Studio software (LI-COR), and assembled for presentation using Photoshop and Illustrator software (Adobe).

### Production of concentrated lentiviral particles

Lentiviruses used for transduction of mouse glia were concentrated after large scale packaging as described previously (*33, 67, 68*). Lenti-X 293T cells (Takara Bio Cat#632180) were plated on 150 mm diameter dishes (∼2.5 million cells / dish) and cultured at 37°C, 5% CO_2_ in DMEM supplemented with 10% FCS, 100 U/ml penicillin, 100 μg/ml streptomycin, and 2 mM L-glutamine. After 16-24 hr, cells were transfected using the polyethylenimine (PEI) method (*67*) at a 2:1 PEI:DNA ratio with the following plasmids (amount of DNA per dish in parenthesis): psPAX2 (18 μg), pMD2.G (11.25 μg), and a plasmid encoding either Gαi bONE-GO WT or Gαi bONE-GO WA (i.e., bearing the W139A mutation in GINIP) biosensor (27 μg). Approximately 16 hr after transfection, media was replaced. Lentivirus containing media was collected 24 and 48 hr after the initial media change (∼70 mL per dish and 4 dishes for each construct). Media was centrifuged for 5 minutes at 900 x g and filtered through a 0.45 μm sterile PES filter (Fisherbrand cat# FB12566505). Filtered media was centrifuged for ∼18 hr at 17,200 x g at 4°C (Sorvall RC6+, ThermoScientific F12-6x500 LEX rotor) to sediment lentiviral particles. Pellets were washed and gently resuspended in 1 mL of PBS and centrifuged at 50,000 x g for 1 hr at 4°C (Beckman Optima MAX-E, TLA-55 rotor). Pellets were resuspended in 300 μl of PBS to obtain concentrated lentiviral stocks that were stored at −80°C in aliquots. Each aliquot was thawed only once and used for less than a week stored at 4°C for subsequent experiments.

### Mouse primary cortical astroglial cell culture

All animal procedures were approved by the Institutional Animal Care and Use Committee (IACUC) at Boston University Chobanian & Avedisian School of Medicine (PROTO202000018). C57BL/6N wild-type mice were from an in-house colony originally established with animals obtained from the Mutant Mouse Resource & Research Centers (MMRRC) at UC Davis. Astrocyte-rich glial cultures were prepared from the cortex of neonatal mice as previously described (*69*) with modifications. Newborn mouse pups (P1-3) were euthanized by decapitation. Brains were removed from the skull and placed in cold HBSS. The cerebrum was detached from other brain regions under a stereomicroscope by removal of the olfactory bulb and cerebellum, and meninges were peeled off with a tweezer. The cortex was dissected out with forceps by removing the hippocampus and the entire midbrain region. The cortex was minced into approximately 1-2 mm pieces using a sterile razor blade, and digested with 0.05% (w:v) trypsin in HBSS for 10 minutes at 37°C. Trypsinized tissue was washed three times with HBSS to remove trypsin, and resuspended in DMEM supplemented with 10% FBS (Gibco cat# 2614-079), 100 U/ml penicillin, 100 μg/ml streptomycin (complete neuro DMEM) before passing through a sterile 40 μm cell strainer (Fisherbrand, cat# 22363547) to obtain a cell suspension. Six-well plates were coated overnight with 0.1 mg/ml poly-L-lysine hydrobromide (Millipore Sigma Cat#P9155), washed three times with HBSS, and approximately 1.5 millions cells were plated in each well. Media was changed the following day, and cells were subsequently split at a 1:2 ratio every 2-3 days by trypsinization followed by centrifugation at 180 x g for 5 minutes before resuspending and reseeding in complete neuro DMEM. Cells were cultured for not more than 5 passages.

### Transduction of mouse astroglial cells with bONE-GO sensor and BRET measurements

bONE-GO biosensors were expressed in astroglial cells by lentiviral transduction as previously described (*33*) using concentrated stocks described in “*Production of concentrated lentiviral particles*”. Mouse astroglial cells were seeded on 5 mm glass coverslips (Word Precision Instruments cat# 502040) precoated with 0.1 mg/mL poly-L-lysine hydrobromide (overnight incubation followed by 3 washes with HBSS) and placed in a 96-well plate (40,000 cells per well). Approximately 18 hr after seeding, cells were transduced by replacing the media with 100 μl of fresh media supplemented with 6 µg/ml polybrene and lentiviruses for the expression of Gαi bONE-GO (1:1000-1:3000 dilution). Plates were spun at 600 x g for 30 minutes and returned to the incubator. Media was replaced ∼24 hr later.

Kinetic BRET recordings were performed ∼48 hr post-transduction as described below. Coverslips were washed with 200 µl BRET buffer and transferred to a well of a white opaque 96-well plate containing BRET buffer and Nano-Glo (final dilution 1:200) with tweezers, followed by incubation in the dark at room temperature for 2 minutes before measuring luminescence in a PHERAstar OMEGA plate reader (BMG Labtech). Luminescence signals at 450 ± 40 and 535 ± 15 nm were measured at 28°C with a signal integration time of 0.96 s. Adenosine was added as indicated in the figures during the recordings using built-in injectors. BRET was calculated as the ratio between the emission intensity at 535 nm divided by the emission intensity at 450 nm, followed by multiplication by 10^3^. Kinetic traces are represented as change in BRET after subtraction of the baseline signal measured for 30 s before GPCR stimulation [ΔBRETꞏ10^3^ (baseline)]. Where indicated in the figures or figure legends, cells expressing Gαi bONE-GO WT were treated overnight with 0.1 μg/ml pertussis toxin (List Biological Labs, cat#179A). For the concentration-response curve presented in **Fig. 7C**, the average [ΔBRETꞏ10^3^ (baseline)] of the “Buffer” condition was first subtracted from all traces, and then the area-under-curve (AUC) was calculated in Prism (GraphPad) for each trace, followed by curve fit to a 3-parameter sigmoidal equation.

## AUTHOR CONTRIBUTIONS

A.L., R.J., J.Z., and C.E.P., conducted experiments. A.L. and M.G-M. designed experiments and analyzed data.

A.L. and M.G-M. wrote the manuscript with input from all authors. M.G-M. conceived and supervised the project.

## ACKNOWLEDGMENTS

This work was primarily supported by NIH grants R01GM147931 and R01NS117101 (to M.G-M.). A.L. was supported by a F31 Ruth L. Kirschstein NRSA Predocotral Fellowship (F31NS115318) and R.J. was supported by a Predoctoral Fellowship from the American Heart Association (898932). We thank the Boston University Flow Cytometry Core Facility for access to instrumentation and technical support. We thank the following investigators for providing DNA plasmids: N. Lambert (Augusta University, Augusta, GA), P. Slessinger (Mount Sinai NY), A. Kovoor (University of Rhode Island), J. Blumer (Medical University of South Carolina).

---

**Figure S1.**
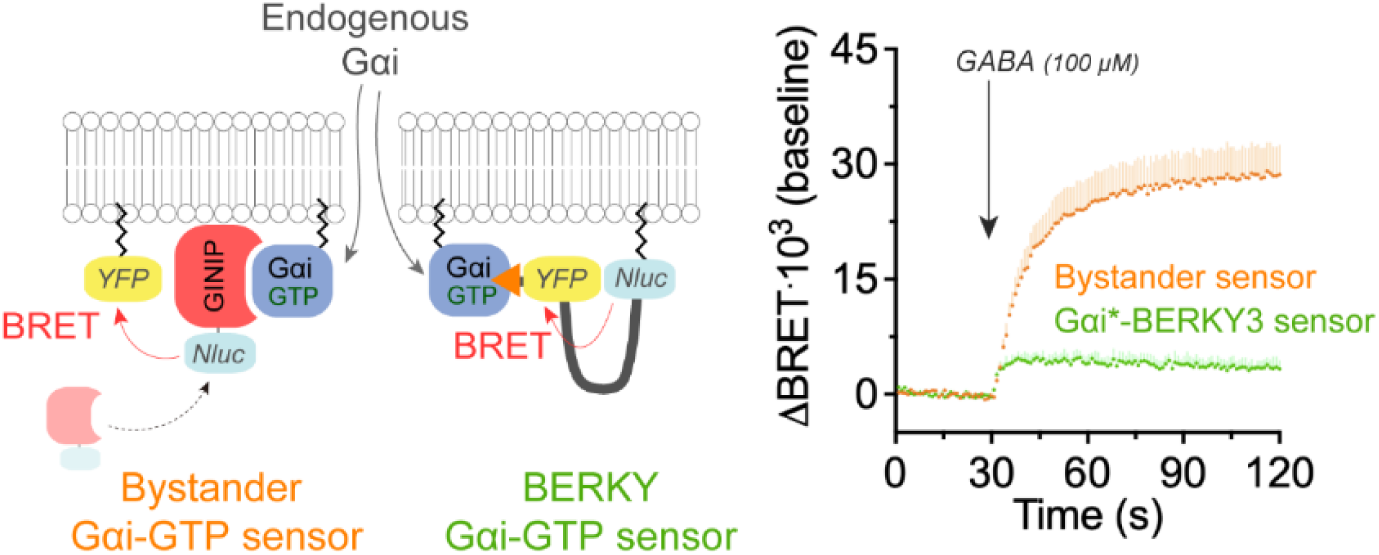
Comparison of bystander Gαi-GTP sensor to Gαi*-BERKY3 sensor for detecting endogenous Gαi. *Left*, Diagram showing the detection of endogenous Gαi-GTP by either Gαi-GTP bystander BRET sensor or BERKY Gαi-GTP sensor. *Right,* Kinetic BRET measurements were carried out in HEK293T cells expressing GABA_B_R and either the Gαi-GTP bystander sensor components (GINIP-Nluc and YFP-CAAX, orange) or Gαi*-BERKY3 sensor (green). Cells were treated with GABA as indicated. Mean ± S.E.M., n=4.

**Figure S2.**
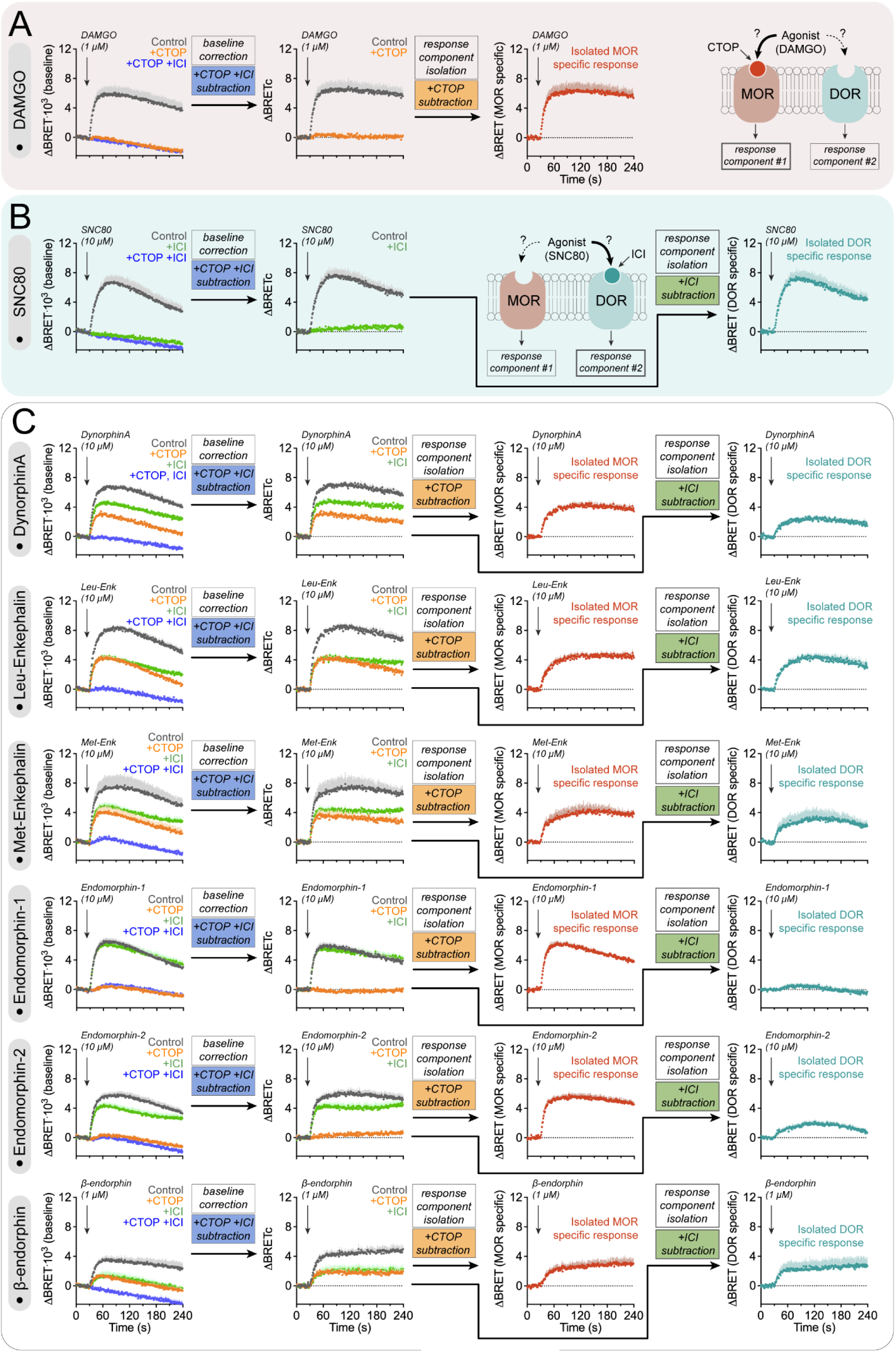
Agonist efficacy of opioid neuropeptides on endogenous receptors in SH-SY5Y. **(A)** Benchmarking of full agonist MOR-specific response in SH-SY5Y cells with the Gαi bONE-GO sensor. Kinetic BRET measurements were carried out in SH-SY5Y cells stably expressing the Gαi bONE-GO sensor in the absence (Control, grey) or presence of 10 μM CTOP (+CTOP, orange), or 10 μM CTOP and 100 μM ICI174,864 (+CTOP +ICI, blue), followed by stimulation with DAMGO. To isolate the MOR-specific response component, first the baseline trace obtained in the presence of CTOP and ICI (+CTOP +ICI) was subtracted from other measurements, followed by the subtraction of the +CTOP trace from the control. Mean ± S.E.M., n=4. **(B)** Benchmarking of full agonist DOR-specific response in SH-SY5Y cells with the Gαi bONE-GO sensor. Kinetic BRET measurements were carried out as in (A) in the absence (Control, grey) or presence of 100 μM ICI174,864 (+ICI, green) or 10 μM CTOP and 100 μM ICI174,864 (+CTOP +ICI, blue), followed by stimulation with SNC80. To isolate the DOR-specific response component, first the baseline trace obtained in the presence of CTOP and ICI (+CTOP +ICI) was subtracted from other measurements, followed by the subtraction of the +ICI trace from the control. Mean ± S.E.M., n=4. **(C)** Isolation of MOR- and DOR-specific responses elicited by opioid neuropeptides in SH-SY5Y cells with Gαi bONE-GO sensor. Kinetic BRET measurements were carried out as in (A) in the absence (Control, grey) or presence of 10 μM CTOP (+CTOP, orange), 100 μM ICI174,864 (+ICI, green), or 10 μM CTOP and 100 μM ICI174,864 (+CTOP +ICI, blue), followed by stimulation with Dynorphin A, Leu-Enkephalin, Met-Enkephalin, Endormorphin-1, Endomorphin-2, or β-endorphin, as indicated. To isolate the MOR- and DOR-specific response components, data were processed as in (A) for the MOR-specific component or as in (B) for the DOR-specific component. Mean ± S.E.M., n=4 (for DAMGO and SNC80), n=4 (for Dynorphin A, Leu-Enkephalin, Met-Enkephalin, and β-endorphin), n=3 (for endomorphin-1, and endomorphin-2). Panels A and B, and the Dynorphin A dataset in panel C contain data also presented in Figure 6.

## REFERENCES

1. A. G. Gilman, G proteins: transducers of receptor-generated signals. Annual review of biochemistry 56, 615–649 (1987)10.1146/annurev.bi.56.070187.003151).

2. J. Marx, Nobel Prizes. Medicine: a signal award for discovering G proteins. *Science (New York*, N.Y 266, 368–369 (1994); published online EpubOct 21 (

3. W. M. Oldham, H. E. Hamm, Heterotrimeric G protein activation by G-protein-coupled receptors. Nature reviews. Molecular cell biology 9, 60–71 (2008); published online EpubJan (10.1038/nrm2299).

4. A. J. Morris, C. C. Malbon, Physiological regulation of G protein-linked signaling. Physiological reviews 79, 1373–1430 (1999); published online EpubOct (

5. W. I. Weis, B. K. Kobilka, The Molecular Basis of G Protein-Coupled Receptor Activation. Annual review of biochemistry 87, 897–919 (2018); published online EpubJun 20 (10.1146/annurev-biochem-060614-033910).

6. J. S. Smith, R. J. Lefkowitz, S. Rajagopal, Biased signalling: from simple switches to allosteric microprocessors. Nature reviews. Drug discovery 17, 243–260 (2018); published online EpubApr (10.1038/nrd.2017.229).

7. A. de Mendoza, A. Sebe-Pedros, I. Ruiz-Trillo, The evolution of the GPCR signaling system in eukaryotes: modularity, conservation, and the transition to metazoan multicellularity. Genome biology and evolution 6, 606–619 (2014); published online EpubMar (10.1093/gbe/evu038).

8. V. Anantharaman, S. Abhiman, R. F. de Souza, L. Aravind, Comparative genomics uncovers novel structural and functional features of the heterotrimeric GTPase signaling system. Gene 475, 63–78 (2011); published online EpubApr 15 (10.1016/j.gene.2010.12.001).

9. D. Urano, A. M. Jones, Heterotrimeric G protein-coupled signaling in plants. Annual review of plant biology 65, 365–384 (2014)10.1146/annurev-arplant-050213-040133).

10. K. Sriram, P. A. Insel, G Protein-Coupled Receptors as Targets for Approved Drugs: How Many Targets and How Many Drugs? Mol Pharmacol 93, 251–258 (2018); published online EpubApr (10.1124/mol.117.111062).

11. A. L. Hopkins, C. R. Groom, The druggable genome. Nat Rev Drug Discov 1, 727–730 (2002); published online EpubSep (10.1038/nrd892).

12. A. S. Hauser, M. M. Attwood, M. Rask-Andersen, H. B. Schioth, D. E. Gloriam, Trends in GPCR drug discovery: new agents, targets and indications. Nature reviews. Drug discovery 16, 829–842 (2017); published online EpubDec (10.1038/nrd.2017.178).

13. D. S. Sauriyal, A. S. Jaggi, N. Singh, Extending pharmacological spectrum of opioids beyond analgesia: multifunctional aspects in different pathophysiological states. Neuropeptides 45, 175–188 (2011); published online EpubJun (10.1016/j.npep.2010.12.004).

14. A. Manglik, H. Lin, D. K. Aryal, J. D. McCorvy, D. Dengler, G. Corder, A. Levit, R. C. Kling, V. Bernat, H. Hubner, X. P. Huang, M. F. Sassano, P. M. Giguere, S. Lober, D. Da, G. Scherrer, B. K. Kobilka, P. Gmeiner, B. L. Roth, B. K. Shoichet, Structure-based discovery of opioid analgesics with reduced side effects. Nature 537, 185–190 (2016); published online EpubSep 8 (10.1038/nature19112).

15. A. Faouzi, H. Wang, S. A. Zaidi, J. F. DiBerto, T. Che, Q. Qu, M. J. Robertson, M. K. Madasu, A. El Daibani, B. R. Varga, T. Zhang, C. Ruiz, S. Liu, J. Xu, K. Appourchaux, S. T. Slocum, S. O. Eans, M. D. Cameron, R. Al-Hasani, Y. X. Pan, B. L. Roth, J. P. McLaughlin, G. Skiniotis, V. Katritch, B. K. Kobilka, S. Majumdar, Structure-based design of bitopic ligands for the µ-opioid receptor. Nature 613, 767–774 (2023); published online EpubJan (10.1038/s41586-022-05588-y).

16. R. Hill, A. Disney, A. Conibear, K. Sutcliffe, W. Dewey, S. Husbands, C. Bailey, E. Kelly, G. Henderson, The novel μ-opioid receptor agonist PZM21 depresses respiration and induces tolerance to antinociception. Br J Pharmacol 175, 2653–2661 (2018); published online EpubJul (10.1111/bph.14224).

17. S. M. DeWire, D. S. Yamashita, D. H. Rominger, G. Liu, C. L. Cowan, T. M. Graczyk, X. T. Chen, P. M. Pitis, D. Gotchev, C. Yuan, M. Koblish, M. W. Lark, J. D. Violin, A G protein-biased ligand at the μ-opioid receptor is potently analgesic with reduced gastrointestinal and respiratory dysfunction compared with morphine. J Pharmacol Exp Ther 344, 708–717 (2013); published online EpubMar (10.1124/jpet.112.201616).

18. C. L. Schmid, N. M. Kennedy, N. C. Ross, K. M. Lovell, Z. Yue, J. Morgenweck, M. D. Cameron, T. D. Bannister, L. M. Bohn, Bias Factor and Therapeutic Window Correlate to Predict Safer Opioid Analgesics. Cell 171, 1165–1175 e1113 (2017); published online EpubNov 16 (10.1016/j.cell.2017.10.035).

19. E. A. Fink, J. Xu, H. Hübner, J. M. Braz, P. Seemann, C. Avet, V. Craik, D. Weikert, M. F. Schmidt, C. M. Webb, N. A. Tolmachova, Y. S. Moroz, X. P. Huang, C. Kalyanaraman, S. Gahbauer, G. Chen, Z. Liu, M. P. Jacobson, J. J. Irwin, M. Bouvier, Y. Du, B. K. Shoichet, A. I. Basbaum, P. Gmeiner, Structure-based discovery of nonopioid analgesics acting through the α(2A)-adrenergic receptor. Science 377, eabn7065 (2022); published online EpubSep 30 (10.1126/science.abn7065).

20. A. S. Hauser, C. Avet, C. Normand, A. Mancini, A. Inoue, M. Bouvier, D. E. Gloriam, Common coupling map advances GPCR-G protein selectivity. eLife 11, (2022); published online EpubMar 18 (10.7554/eLife.74107).

21. B. Bettler, K. Kaupmann, J. Mosbacher, M. Gassmann, Molecular structure and physiological functions of GABA(B) receptors. Physiological reviews 84, 835–867 (2004); published online EpubJul (10.1152/physrev.00036.2003).

22. B. Paul, S. Sribhashyam, S. Majumdar, Opioid signaling and design of analgesics. Prog Mol Biol Transl Sci 195, 153–176 (2023)10.1016/bs.pmbts.2022.06.017).

23. S. M. Bernhard, J. Han, T. Che, GPCR-G protein selectivity revealed by structural pharmacology. Febs j, (2023); published online EpubDec 27 (10.1111/febs.17049).

24. A. Koehl, H. Hu, S. Maeda, Y. Zhang, Q. Qu, J. M. Paggi, N. R. Latorraca, D. Hilger, R. Dawson, H. Matile, G. F. X. Schertler, S. Granier, W. I. Weis, R. O. Dror, A. Manglik, G. Skiniotis, B. K. Kobilka, Structure of the µ-opioid receptor-G(i) protein complex. Nature 558, 547–552 (2018); published online EpubJun (10.1038/s41586-018-0219-7).

25. M. Bunemann, M. Frank, M. J. Lohse, Gi protein activation in intact cells involves subunit rearrangement rather than dissociation. Proceedings of the National Academy of Sciences of the United States of America 100, 16077–16082 (2003); published online EpubDec 23 (10.1073/pnas.2536719100).

26. C. Gales, J. J. Van Durm, S. Schaak, S. Pontier, Y. Percherancier, M. Audet, H. Paris, M. Bouvier, Probing the activation-promoted structural rearrangements in preassembled receptor-G protein complexes. Nature structural & molecular biology 13, 778–786 (2006); published online EpubSep (10.1038/nsmb1134).

27. B. Hollins, S. Kuravi, G. J. Digby, N. A. Lambert, The c-terminus of GRK3 indicates rapid dissociation of G protein heterotrimers. Cellular signalling 21, 1015–1021 (2009); published online EpubJun (10.1016/j.cellsig.2009.02.017).

28. I. Masuho, O. Ostrovskaya, G. M. Kramer, C. D. Jones, K. Xie, K. A. Martemyanov, Distinct profiles of functional discrimination among G proteins determine the actions of G protein-coupled receptors. Science signaling 8, ra123 (2015); published online EpubDec 1 (10.1126/scisignal.aab4068).

29. C. Janetopoulos, T. Jin, P. Devreotes, Receptor-mediated activation of heterotrimeric G-proteins in living cells. Science 291, 2408–2411 (2001); published online EpubMar 23 (10.1126/science.1055835).

30. R. H. J. Olsen, J. F. DiBerto, J. G. English, A. M. Glaudin, B. E. Krumm, S. T. Slocum, T. Che, A. C. Gavin, J. D. McCorvy, B. L. Roth, R. T. Strachan, TRUPATH, an open-source biosensor platform for interrogating the GPCR transducerome. Nat Chem Biol 16, 841–849 (2020); published online EpubAug (10.1038/s41589-020-0535-8).

31. C. Xu, Y. Zhou, Y. Liu, L. Lin, P. Liu, X. Wang, Z. Xu, J. P. Pin, P. Rondard, J. Liu, Specific pharmacological and G(i/o) protein responses of some native GPCRs in neurons. Nature communications 15, 1990 (2024); published online EpubMar 5 (10.1038/s41467-024-46177-z).

32. M. Maziarz, J. C. Park, A. Leyme, A. Marivin, A. Garcia-Lopez, P. P. Patel, M. Garcia-Marcos, Revealing the Activity of Trimeric G-proteins in Live Cells with a Versatile Biosensor Design. Cell 182, 770–785.e716 (2020); published online EpubAug 6 (10.1016/j.cell.2020.06.020).

33. R. Janicot, M. Maziarz, J. C. Park, J. Zhao, A. Luebbers, E. Green, C. E. Philibert, H. Zhang, M. D. Layne, J. C. Wu, M. Garcia-Marcos, Direct interrogation of context-dependent GPCR activity with a universal biosensor platform. Cell, (2024); published online EpubFeb 16 (10.1016/j.cell.2024.01.028).

34. C. Avet, A. Mancini, B. Breton, C. Le Gouill, A. S. Hauser, C. Normand, H. Kobayashi, F. Gross, M. Hogue, V. Lukasheva, S. St-Onge, M. Carrier, M. Héroux, S. Morissette, E. B. Fauman, J.-P. Fortin, S. Schann, X. Leroy, D. E. Gloriam, M. Bouvier, Effector membrane translocation biosensors reveal G protein and βarrestin coupling profiles of 100 therapeutically relevant GPCRs. eLife 11, e74101 (2022); published online Epub2022/03/18 (10.7554/eLife.74101).

35. F. S. Willard, A. B. Low, C. R. McCudden, D. P. Siderovski, Differential G-alpha interaction capacities of the GoLoco motifs in Rap GTPase activating proteins. Cellular signalling 19, 428–438 (2007); published online Epub2007/02/01/ (10.1016/j.cellsig.2006.07.013).

36. C. K. Webb, C. R. McCudden, F. S. Willard, R. J. Kimple, D. P. Siderovski, G. S. Oxford, D2 dopamine receptor activation of potassium channels is selectively decoupled by Galpha-specific GoLoco motif peptides. J Neurochem 92, 1408–1418 (2005); published online EpubMar (10.1111/j.1471-4159.2004.02997.x).

37. S. Kuravi, T. H. Lan, A. Barik, N. A. Lambert, Third-party bioluminescence resonance energy transfer indicates constitutive association of membrane proteins: application to class a g-protein-coupled receptors and g-proteins. Biophys J 98, 2391–2399 (2010); published online EpubMay 19 (10.1016/j.bpj.2010.02.004).

38. B. R. Martin, N. A. Lambert, Activated G Protein Gαs Samples Multiple Endomembrane Compartments. The Journal of biological chemistry 291, 20295–20302 (2016); published online EpubSep 23 (10.1074/jbc.M116.729731).

39. J. C. Park, A. Luebbers, M. Dao, A. Semeano, A. M. Nguyen, M. P. Papakonstantinou, S. Broselid, H. Yano, K. A. Martemyanov, M. Garcia-Marcos, Fine-tuning GPCR-mediated neuromodulation by biasing signaling through different G protein subunits. Mol Cell 83, 2540–2558.e2512 (2023); published online EpubJul 20 (10.1016/j.molcel.2023.06.006).

40. A. Luebbers, A. J. Gonzalez-Hernandez, M. Zhou, S. J. Eyles, J. Levitz, M. Garcia-Marcos, Dissecting the molecular basis for the modulation of neurotransmitter GPCR signaling by GINIP. Structure 32, 47–59.e47 (2024); published online EpubJan 4 (10.1016/j.str.2023.10.010).

41. S. Gaillard, L. Lo Re, A. Mantilleri, R. Hepp, L. Urien, P. Malapert, S. Alonso, M. Deage, C. Kambrun, M. Landry, S. A. Low, A. Alloui, B. Lambolez, G. Scherrer, Y. Le Feuvre, E. Bourinet, A. Moqrich, GINIP, a Gαi-interacting protein, functions as a key modulator of peripheral GABAB receptor-mediated analgesia. Neuron 84, 123–136 (2014); published online EpubOct 1 (10.1016/j.neuron.2014.08.056).

42. B. F. Binkowski, B. L. Butler, P. F. Stecha, C. T. Eggers, P. Otto, K. Zimmerman, G. Vidugiris, M. G. Wood, L. P. Encell, F. Fan, K. V. Wood, A Luminescent Biosensor with Increased Dynamic Range for Intracellular cAMP. ACS Chemical Biology 6, 1193–1197 (2011); published online Epub2011/11/18 (10.1021/cb200248h).

43. E. S. Levitt, L. C. Purington, J. R. Traynor, Gi/o-coupled receptors compete for signaling to adenylyl cyclase in SH-SY5Y cells and reduce opioid-mediated cAMP overshoot. Molecular pharmacology 79, 461–471 (2011); published online EpubMar (10.1124/mol.110.064816).

44. S. P. H. Alexander, A. Christopoulos, A. P. Davenport, E. Kelly, A. A. Mathie, J. A. Peters, E. L. Veale, J. F. Armstrong, E. Faccenda, S. D. Harding, J. A. Davies, M. P. Abbracchio, G. Abraham, A. Agoulnik, W. Alexander, K. Al-Hosaini, M. Bäck, J. G. Baker, N. M. Barnes, R. Bathgate, J. M. Beaulieu, A. G. Beck-Sickinger, M. Behrens, K. E. Bernstein, B. Bettler, N. J. M. Birdsall, V. Blaho, F. Boulay, C. Bousquet, H. Bräuner-Osborne, G. Burnstock, G. Caló, J. P. Castaño, K. J. Catt, S. Ceruti, P. Chazot, N. Chiang, B. Chini, J. Chun, A. Cianciulli, O. Civelli, L. H. Clapp, R. Couture, H. M. Cox, Z. Csaba, C. Dahlgren, G. Dent, S. D. Douglas, P. Dournaud, S. Eguchi, E. Escher, E. J. Filardo, T. Fong, M. Fumagalli, R. R. Gainetdinov, M. L. Garelja, M. de Gasparo, C. Gerard, M. Gershengorn, F. Gobeil, T. L. Goodfriend, C. Goudet, L. Grätz, K. J. Gregory, A. L. Gundlach, J. Hamann, J. Hanson, R. L. Hauger, D. L. Hay, A. Heinemann, D. Herr, M. D. Hollenberg, N. D. Holliday, M. Horiuchi, D. Hoyer, L. Hunyady, A. Husain, I. J. AP, T. Inagami, K. A. Jacobson, R. T. Jensen, R. Jockers, D. Jonnalagadda, S. Karnik, K. Kaupmann, J. Kemp, C. Kennedy, Y. Kihara, T. Kitazawa, P. Kozielewicz, H. J. Kreienkamp, J. P. Kukkonen, T. Langenhan, D. Larhammar, K. Leach, D. Lecca, J. D. Lee, S. E. Leeman, J. Leprince, X. X. Li, S. J. Lolait, A. Lupp, R. Macrae, J. Maguire, D. Malfacini, J. Mazella, C. A. McArdle, S. Melmed, M. C. Michel, L. J. Miller, V. Mitolo, B. Mouillac, C. E. Müller, P. M. Murphy, J. L. Nahon, T. Ngo, X. Norel, D. Nyimanu, A. M. O’Carroll, S. Offermanns, M. A. Panaro, M. Parmentier, R. G. Pertwee, J. P. Pin, E. R. Prossnitz, M. Quinn, R. Ramachandran, M. Ray, R. K. Reinscheid, P. Rondard, G. E. Rovati, C. Ruzza, G. J. Sanger, T. Schöneberg, G. Schulte, S. Schulz, D. L. Segaloff, C. N. Serhan, K. D. Singh, C. M. Smith, L. A. Stoddart, Y. Sugimoto, R. Summers, V. P. Tan, D. Thal, W. W. Thomas, P. Timmermans, K. Tirupula, L. Toll, G. Tulipano, H. Unal, T. Unger, C. Valant, P. Vanderheyden, D. Vaudry, H. Vaudry, J. P. Vilardaga, C. S. Walker, J. M. Wang, D. T. Ward, H. J. Wester, G. B. Willars, T. L. Williams, T. M. Woodruff, C. Yao, R. D. Ye, The Concise Guide to PHARMACOLOGY 2023/24: G protein-coupled receptors. Br J Pharmacol 180 **Suppl 2**, S23–s144 (2023); published online EpubOct (10.1111/bph.16177).

45. K. Raynor, H. Kong, Y. Chen, K. Yasuda, L. Yu, G. I. Bell, T. Reisine, Pharmacological characterization of the cloned kappa-, delta-, and mu-opioid receptors. Molecular pharmacology 45, 330–334 (1994); published online EpubFeb (

46. J. Gong, J. A. Strong, S. Zhang, X. Yue, R. N. DeHaven, J. D. Daubert, J. A. Cassel, G. Yu, E. Mansson, L. Yu, Endomorphins fully activate a cloned human mu opioid receptor. FEBS letters 439, 152–156 (1998); published online EpubNov 13 (10.1016/s0014-5793(98)01362-3).

47. R. B. Dias, D. M. Rombo, J. A. Ribeiro, J. M. Henley, A. M. Sebastião, Adenosine: setting the stage for plasticity. Trends Neurosci 36, 248–257 (2013); published online EpubApr (10.1016/j.tins.2012.12.003).

48. T. Kenakin, Is the Quest for Signaling Bias Worth the Effort? Molecular pharmacology 93, 266–269 (2018); published online EpubApr (10.1124/mol.117.111187).

49. T. Kenakin, Biased Receptor Signaling in Drug Discovery. Pharmacological reviews 71, 267–315 (2019); published online EpubApr (10.1124/pr.118.016790).

50. T. Che, H. Dwivedi-Agnihotri, A. K. Shukla, B. L. Roth, Biased ligands at opioid receptors: Current status and future directions. Science signaling 14, (2021); published online EpubApr 6 (10.1126/scisignal.aav0320).

51. L. M. Bohn, R. J. Lefkowitz, R. R. Gainetdinov, K. Peppel, M. G. Caron, F. T. Lin, Enhanced morphine analgesia in mice lacking beta-arrestin 2. Science 286, 2495–2498 (1999); published online EpubDec 24 (10.1126/science.286.5449.2495).

52. A. Kliewer, A. Gillis, R. Hill, F. Schmiedel, C. Bailey, E. Kelly, G. Henderson, M. J. Christie, S. Schulz, Morphine-induced respiratory depression is independent of β-arrestin2 signalling. Br J Pharmacol 177, 2923–2931 (2020); published online EpubJul (10.1111/bph.15004).

53. A. Gillis, A. B. Gondin, A. Kliewer, J. Sanchez, H. D. Lim, C. Alamein, P. Manandhar, M. Santiago, S. Fritzwanker, F. Schmiedel, T. A. Katte, T. Reekie, N. L. Grimsey, M. Kassiou, B. Kellam, C. Krasel, M. L. Halls, M. Connor, J. R. Lane, S. Schulz, M. J. Christie, M. Canals, Low intrinsic efficacy for G protein activation can explain the improved side effect profiles of new opioid agonists. Science signaling 13, (2020); published online EpubMar 31 (10.1126/scisignal.aaz3140).

54. A. Gillis, A. Kliewer, E. Kelly, G. Henderson, M. J. Christie, S. Schulz, M. Canals, Critical Assessment of G Protein-Biased Agonism at the μ-Opioid Receptor. Trends Pharmacol Sci 41, 947–959 (2020); published online EpubDec (10.1016/j.tips.2020.09.009).

55. T. H. Lan, Q. Liu, C. Li, G. Wu, N. A. Lambert, Sensitive and high resolution localization and tracking of membrane proteins in live cells with BRET. Traffic 13, 1450–1456 (2012); published online EpubNov (10.1111/j.1600-0854.2012.01401.x).

56. Q. Wan, N. Okashah, A. Inoue, R. Nehmé, B. Carpenter, C. G. Tate, N. A. Lambert, Mini G protein probes for active G protein-coupled receptors (GPCRs) in live cells. The Journal of biological chemistry 293, 7466–7473 (2018); published online EpubMay 11 (10.1074/jbc.RA118.001975).

57. N. G. Tsvetanova, M. Trester-Zedlitz, B. W. Newton, G. E. Peng, J. R. Johnson, D. Jimenez-Morales, A. P. Kurland, N. J. Krogan, M. von Zastrow, Endosomal cAMP production broadly impacts the cellular phosphoproteome. The Journal of biological chemistry 297, 100907 (2021); published online EpubJul (10.1016/j.jbc.2021.100907).

58. A. Radoux-Mergault, L. Oberhauser, S. Aureli, F. L. Gervasio, M. Stoeber, Subcellular location defines GPCR signal transduction. Sci Adv 9, eadf6059 (2023); published online EpubApr 21 (10.1126/sciadv.adf6059).

59. M. Stoeber, D. Jullié, B. T. Lobingier, T. Laeremans, J. Steyaert, P. W. Schiller, A. Manglik, M. von Zastrow, A Genetically Encoded Biosensor Reveals Location Bias of Opioid Drug Action. Neuron 98, 963–976.e965 (2018); published online EpubJun 6 (10.1016/j.neuron.2018.04.021).

60. G. A. Kumar, M. A. Puthenveedu, Diversity and specificity in location-based signaling outputs of neuronal GPCRs. Curr Opin Neurobiol 76, 102601 (2022); published online EpubOct (10.1016/j.conb.2022.102601).

61. C. S. Kearn, K. Blake-Palmer, E. Daniel, K. Mackie, M. Glass, Concurrent stimulation of cannabinoid CB1 and dopamine D2 receptors enhances heterodimer formation: a mechanism for receptor cross-talk? Molecular pharmacology 67, 1697–1704 (2005); published online EpubMay (10.1124/mol.104.006882).

62. S. S. Oner, N. An, A. Vural, B. Breton, M. Bouvier, J. B. Blumer, S. M. Lanier, Regulation of the AGS3ꞏG{alpha}i signaling complex by a seven-transmembrane span receptor. The Journal of biological chemistry 285, 33949–33958 (2010); published online EpubOct 29 (10.1074/jbc.M110.138073).

63. Y. Tang, L. A. Hu, W. E. Miller, N. Ringstad, R. A. Hall, J. A. Pitcher, P. DeCamilli, R. J. Lefkowitz, Identification of the endophilins (SH3p4/p8/p13) as novel binding partners for the beta1-adrenergic receptor. Proceedings of the National Academy of Sciences of the United States of America 96, 12559–12564 (1999); published online EpubOct 26 (10.1073/pnas.96.22.12559).

64. K. Ishii, L. Hein, B. Kobilka, S. R. Coughlin, Kinetics of thrombin receptor cleavage on intact cells. Relation to signaling. The Journal of biological chemistry 268, 9780–9786 (1993); published online EpubMay 5 (

65. M. Maziarz, J. C. Park, A. Leyme, A. Marivin, A. Garcia-Lopez, P. P. Patel, M. Garcia-Marcos, Revealing the Activity of Trimeric G-proteins in Live Cells with a Versatile Biosensor Design. Cell 182, 770–785 e716 (2020); published online EpubAug 6 (10.1016/j.cell.2020.06.020).

66. Y. A. Bochkov, A. C. Palmenberg, Translational efficiency of EMCV IRES in bicistronic vectors is dependent upon IRES sequence and gene location. BioTechniques 41, 283–284, 286, 288 passim (2006); published online EpubSep (10.2144/000112243).

67. P. A. Longo, J. M. Kavran, M. S. Kim, D. J. Leahy, Transient mammalian cell transfection with polyethylenimine (PEI). Methods Enzymol 529, 227–240 (2013)10.1016/b978-0-12-418687-3.00018-5).

68. R. Janicot, J. C. Park, M. Garcia-Marcos, Detecting GPCR Signals With Optical Biosensors of Galpha-GTP in Cell Lines and Primary Cell Cultures. Current protocols 3, e796 (2023); published online EpubJun (10.1002/cpz1.796).

69. S. Kaech, G. Banker, Culturing hippocampal neurons. Nat Protoc 1, 2406–2415 (2006)10.1038/nprot.2006.356).

